# Andean uplift, drainage basin formation, and the evolution of riverweeds *(Marathrum,* Podostemaceae) in northern South America

**DOI:** 10.1101/2020.11.13.382200

**Authors:** Ana M. Bedoya, Adam D. Leaché, Richard G. Olmstead

**Affiliations:** Department of Biology and Burke Museum, University of Washington, Seattle, WA, United States

**Keywords:** Andean uplift, Marathrum, next-generation sequencing, northern South America, phylogenomics, rivers, target enrichment

## Abstract

- Northern South America is a geologically dynamic and species-rich region. While fossil and stratigraphic data show that reconfiguration of river drainages resulted from mountain uplift in the tropical Andes, investigations of the impact of landscape change on the evolution of the flora in the region have been restricted to terrestrial taxa.
- We explore the role of landscape change on the evolution of plants living strictly in rivers across drainage basins in northern South America by conducting population structure, phylogenomic, phylogenetic networks, and divergence-dating analyses for populations of riverweeds (*Marathrum*, Podostemaceae).
- We show that mountain uplift and drainage basin formation isolated populations of *Marathrum* and created barriers to gene flow across rivers drainages. Sympatric species hybridize and the hybrids show the phenotype of one parental line. We propose that the pattern of divergence of populations reflect the formation of river drainages, which was not complete until <4 Ma
- Our study provides a clear picture of the role of landscape change in shaping the evolution of riverweeds in northern South America, advances our understanding of the reproductive biology of this remarkable group of plants, and spotlights the impact of hybridization in phylogenetic inference.

## Introduction

Landscape change has been recognized as a major driver of diversification of lineages (Hoorn *et al.*, 2013). In the Neotropics, the most species-rich region in the world, the uplift of the Andean Cordillera is thought to have contributed directly and indirectly to the assembly of the terrestrial biota (Janzen, 1967; Kattan *et al.*, 2004; Antonelli & Sanmartín, 2011; Sklenář *et al.*, 2011; Smith *et al.*, 2014; Hoorn *et al.*, 2018; Quintero & Jetz, 2018). However, by shifting the physical locations of watershed boundaries, connecting and disconnecting adjacent river basins, (Albert & Crampton, 2010; Hoorn *et al.*, 2010; Tagliacollo *et al.*, 2015; Ruokolainen *et al.*, 2019) the uplift of the Andean Cordillera has also impacted the evolution of organisms living in rivers as has been shown in fish (Albert *et al.*, 2006, 2020; Picq *et al.*, 2014; Tagliacollo *et al.*, 2015). The role of Andean uplift in Neotropical plant evolution has been investigated in terrestrial groups living in mountains (Richardson *et al.*, 2018), where a correlation of Andean orogeny with increased diversification rates (Lagomarsino *et al.*, 2016; Pérez-Escobar *et al.*, 2017; Testo *et al.*, 2019), and explosive radiation in taxa in high-elevation tropical ecosystems (Hughes & Eastwood, 2006; Madriñán *et al.*, 2013; Nürk *et al.*, 2013; Nevado *et al.*, 2018) have been inferred. However, the impact of Andean uplift and landscape change in the region on plants in rivers remains unaddressed.

Here, we investigate the role of drainage basin formation linked to Andean uplift in shaping the evolution of river plants using riverweeds in the genus *Marathrum* (Podostemaceae) as a model system. Unlike most other angiosperms, except for the Hydrostachyaceae which are restricted to Africa and Madagascar, riverweeds are herbs that live attached to rocks partially or completely submerged in fast-flowing water ecosystems like river-rapids and waterfalls (Fig. **1b**) (van Royen, 1951; Philbrick & Novelo, 1995). The family has a Pantropical distribution and is the only group of aquatic plants in the Neotropics to live strictly in rivers (van Royen, 1951; Tippery *et al.*, 2011). Riverweeds are mostly annuals with short generation times and have flowers with a reduced perianth and capsular fruits that are produced and project above the water surface in the dry season (Willis, 1915; Philbrick & Novelo, 1998; Koi *et al.*, 2015). Hundreds of seeds are shed prior to the onset of the wet season when the water level is still low, and attach to the rock with a sticky mucilage (Reyes-Ortega *et al.*, 2009). The adult plants remain vegetative and attached to rocks via adhesive hairs and a biofilm of cyanobacteria (Rutishauser *et al.*, 1999; Jäger-Zürn & Grubert, 2000). Therefore, it has been suggested that the Podostemaceae have a reduced dispersal ability and are highly endemic (Philbrick *et al.*, 2010).

**Figure 1.**
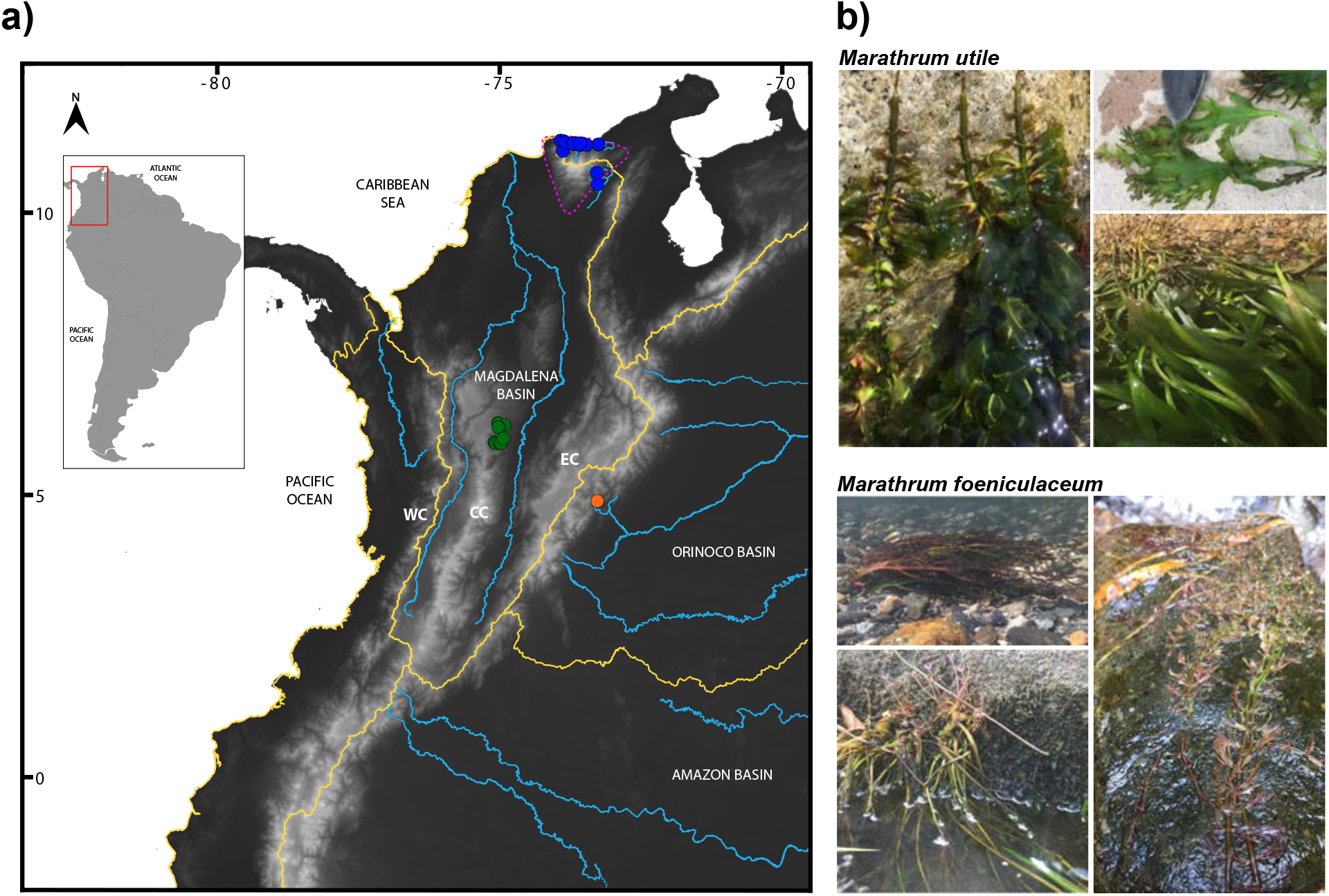
a) Localities where *Marathrum* specimens were sampled (dots). Green=Antioquia, Blue=Caribe, Orange=Boyacá. The three Andean Cordilleras are shown. WC= Western Cordillera, CC=Central cordillera, EC=Eastern Cordillera. Yellow lines delimit drainage basins. The Sierra Nevada de Santa Marta is indicated with pink dashed lines. Major rivers are shown in blue. b) Phenotype of *M. utile* and *M. foeniculaceum*. The former one has laminar, entire leaves. The latter has highly dissected leaves. Flower morphology varies geographically in both species.

Within the Podostemaceae, the genus *Marathrum* has 22 recognized species (van Royen, 1951; Novelo & Philbrick, 1997; Novelo *et al.*, 2009; Tippery *et al.*, 2011). The genus is distributed in the West Indies, Central America, and across the Andes in northern South America (nSA) (van Royen, 1951), making it the ideal model system for this study. The taxonomic classification of the group is confounded by the high degree of modification of vegetative organs (Rutishauser, 1995, 1997; Rutishauser *et al.*, 1999), as well as the reduced number of reproductive characters useful for classification (Philbrick, pers. comm.), which has resulted in multiple species proposed to be synonyms (Novelo *et al.*, 2009; Tippery *et al.*, 2011). Two species in particular are distinct from one another in their leaf type; *Marathrum utile* Tul. is the only species in the group with entire, laminar leaves (Fig. **1b**) (van Royen, 1951), whereas *Marathrum foeniculaceum* Bonpl. has highly dissected leaves (Fig. **1b**) and includes several other recognized species that have been proposed to be synonyms to it (Novelo *et al.*, 2009; Tippery *et al.*, 2011).

Palynological, stratigraphic, and macrofossil data (Latrubesse *et al.*, 2010; Hoorn *et al.*, 2010) support a landscape evolution model where from the Late Miocene (*ca.* 11Ma) to present, exceptionally rapid uplift of the Andes transformed the aquatic ecosystems in the region by causing a shift from a wetland-dominated environment, *Pebas Lake*, to the formation of current drainage basins interrupted by the Andes (*i.e.* Pacific, Magdalena, Orinoco, and Amazon; Fig. **1a**. Important features in the landscape of nSA include the Andean Cordillera, which in the region is divided into three ranges, *i.e.* the Western Cordillera (WC), Central Cordillera (CC), and Eastern Cordillera (EC) (Fig. **1a**). Another important topographic element is the Sierra Nevada de Santa Marta (SNSM) (Fig. **1a**), which is a separate formation from the Andes and is the highest coastal relief on earth (Duque-Caro, 1979; Villagómez *et al.*, 2011; Parra *et al.*, 2019). Multiple rivers originate in the SNSM (Fig. **1b**), with evidence for rapid and recent uplift of this mountain system (Villagómez *et al.*, 2011; Parra *et al.*, 2019). The rivers on the southern slopes of the SNSM are part of the Magdalena drainage basin (Abell *et al.*, 2008). However, these rivers are not connected through fast-flowing water currents to the other tributaries of the Magdalena drainage basin.

Given the dynamic nature of the landscape in northern South America and its role in shaping rivers, in this study we investigate how landscape change shaped the evolution of *Marathrum* in the region. We generate target enrichment and ddRADseq data and infer the genetic constitution of populations of *Marathrum* located across the Andes and the SNSM. We determine if there is gene flow across drainage basins and reconstruct the evolutionary relationships of populations of *Marathrum* using data concatenation, species tree, and phylogenetic network approaches. Using a species tree and a time-calibrated tree, we provide a hypothesis for the pattern and timing of divergence of populations of *Marathrum* currently disconnected by the Andes, the SNSM, and inter-Andean valleys. We propose that this pattern reflects the evolution of drainage basins in northern South America. Furthermore, this study provides an insight into the reproductive biology of *Marathrum* and shows that hybridization confounds phylogenetic inference of its populations.

## Materials and Methods

### Data collection, library preparation and sequencing

We collected 118 individuals; 115 of *Marathrum* (75 individuals of *M. foeniculaceum* and 40 individuals of *M. utile*. See Fig. **1b**) and three of *Weddellina squamulosa* Tul. from Antioquia, (eastern slope of the CC), Boyacá (eastern slope of the EC), and Caribe (SNSM) (Fig. **1a**). These populations are currently disconnected by the Andes, the SNSM, inter-Andean valleys, and the lowlands between the Andes and the SNSM (Fig. **1a**). Samples were preserved as vouchers, and tissues were taken from leaf material and stored in silica gel. Voucher information is shown in Table S1. We extracted DNA from silica-dried leaf material using a modified CTAB protocol (Doyle & Doyle, 1987) and purified the DNA extracts with isopropanol precipitation. We assessed DNA concentration using a Qubit fluorometer and DNA fragmentation levels using gel electrophoresis (1% agarose gels).

#### Double digest Restriction Associated DNA sequencing (ddRADseq)

We prepared ddRADseq libraries for 115 samples following the protocol described by Peterson *et al.*, 2012. In brief, we used *PstI* and *MspI* to double digest *ca.* 500 ng high-molecular DNA and purified fragments with Sera-Mag SpeedBeads prior to ligation of barcoded Illumina adapters. Libraries were size selected to 415–515 bp on a Pippin prep (Sage Science) using a modified protocol where pooled samples were each run in a separate lane and size selected in reference to one lane running only the R2 marker reagent. Final library amplification used Illumina indexed primers and HF-Phusion TAQ. We determined library size distributions and concentrations on an Agilent 2200 TapeStation. Samples were sent to the QB3 sequencing facility at UC Berkeley, where qPCR was performed to determine sequenceable library concentrations prior to multiplexing equimolar amounts of each library and sequencing 150 single-end reads in two lanes of an Illumina HiSeq 4000.

#### Low-depth reference genome of Marathrum

To identify single-copy orthologs for phylogenomic analyses, we collected genome-skimming data for one individual of *Marathrum utile* (Table S1) generated at the QB3 sequencing facility (Berkeley, CA). Total DNA was sheared to a target size of 300–500 bp and 150 paired-end reads were sequenced in an Illumina HiSeq 4000. In order to select a set of target sequences that are nuclear, single-copy, and have orthologs across *Marathrum*, we applied the methods in Chau *et al.*, 2018. Briefly, the *de novo* assembled genome of *Marathrum utile*, a transcriptome of *Hypericum perforatum* L. (Hypericaceae) (assembly generated by the 1kp project, sample code No. BNDE) (One Thousand Plant Transcriptomes Initiative, 2019; Carpenter *et al.*, 2019), a plastome of *Garcinia mangostana* L. (Calophyllaceae), and a mitochondriome of *Populus tremula* L. (Salicaceae) (see Table S1 for sample information and accession numbers) were used as input for the pipeline Sondovac (Schmickl et al., 2016). Sondovac removes reads that map to the plastome or mitochondriome, and duplicated transcripts from the transcriptome and finds genome reads matching the remaining unique transcripts, which are *de novo* assembled. The pipeline filters the targets by length (minimum 600 bp for all contigs for a transcript, and 120 bp for each contig) and uniqueness.

We complemented the set with the APVO SSC and pentatricopeptide repeat genes, which are shared across angiosperms and have been found to be useful for resolution of inter-specific relationships in some plant groups (Yuan et al., 2009, 2010; Duarte et al., 2010). The sequences were downloaded from http://www.arabidopsis.org and used to conduct searches against the *M. utile* contigs in the draft genome with BLASTN. Up to five of the top hits with a bit score >70 were retained. When hits for the same loci overlapped from different contigs, we assembled the contigs using *de novo* assembly in Geneious v9.1.8. We removed all hits matching the same locus with <95% pairwise identity as they could represent paralogs. We retained individual sequences of at least 120 bp, and loci with at least 600 bp length.

#### Target enrichment

Taxon sampling for the target enrichment experiment was guided by a phylogenetic analysis of all the ddRADseq data using SVDQuartets (Fig. S1) (Chifman & Kubatko, 2014). Of the 118 samples collected, we selected 46 individuals for target enrichment with the aims of 1) including specimens with high-quality genomic DNA and high coverage across the ddRADseq assembly, 2) sampling multiple individuals per populations in each drainage basin, 3) providing equal representation of the clades identified by the ddRADseq data. Probe design and sequencing were performed by RapidGenomics (Gainesville, Florida, US). Briefly, 250–1000 ng of high-molecular DNA was fragmented to a target size of 400 bp. Enriched pools were combined in equimolar ratios and probes were hybridized to pools to enrich for targets. Probes targeting a loci space of 214,484 bp for 536 exons in 218 loci with 2x tiling density were designed and 150 paired-end reads were sequenced in 25% of a lane of an Illumina HiSeq 3000. All raw sequence reads from the low-coverage genome, target enrichment experiment, and ddRADseq datasets are deposited in the National Center for Biotechnology Information (NCBI) Sequence Read Archive (Table S1).

### Data processing

#### ddRADseq

Following the exclusion of 20 samples that failed sequencing or had low coverage, ddRADseq data for the remaining 95 samples were quality checked with FastQC (https://www.bioinformatics.babraham.ac.uk/projects/fastqc/) and demultiplexed and assembled in ipyrad v0.9.42 (Eaton & Overcast, 2020) using a clustering threshold of 0.88. We removed consensus sequences that had low coverage (6 reads), excessive undetermined or heterozygous sites (>12), an excess of shared heterozygosity among samples (paralog filter=0.5), and included loci present in at least four individuals. Preliminary inspection of our resulting assemblies showed that there where high levels of missing data in our ddRADseq dataset with no loci recovered that are shared across ≥90% of the samples and only 70 loci shared across half of the samples (Table S2). In order to increase the number of shared loci across samples and allow for direct comparison of our two genomic datasets, we selected the ddRADseq samples that were also successfully sequenced in our target enrichment dataset (n=34). Reads from this subset where assembled in ipyrad v0.9.42 (Eaton & Overcast, 2020) with parameters specified as described above for the complete ddRADseq dataset, and we generated two alignments that included loci that were present in at least 17 individuals (*min17*; 50% missing data) and at least 4 individuals (*min4*; ~88% missing data).

#### Low-depth genome

Genome-skimming reads were loaded in FastQC to assess sequence quality. We used Trimmomatic v0.39 (Bolger *et al.*, 2014) to remove adapter sequences. *De novo* assembly was performed with the CLC Genomics Workbench 8 (https://digitalinsights.qiagen.com).

#### Target enrichment

We quality checked the demultiplexed target enrichment reads with FastQC and removed adapter sequences with Trimmomatic. Assembly of loci into genotypes with ambiguity codes may affect the accuracy of phylogenetic inference under the multispecies coalescent model (Andermann *et al.*, 2018). We assembled our target enrichment dataset in order to obtain alleles and not genotypes with ambiguity codes. Resulting paired and unpaired reads were assembled with the pipeline HybPiper to align, assemble mapped reads for each target sequence, and extract the coding sequence from the assembled contigs respectively. The best full-length contig is chosen by Hybpiper based on sequencing depth and percent identity with the reference sequence. Hence, for each individual, locus assembly with Hybpiper results in a single contig (allele) (Johnson *et al.*, 2016). Kates *et al.*, 2018 provided a framework for retrieving phased alleles from target enrichment data. However, the pipeline is appropriate for long-read data and when used in our data, it yielded inaccurate assemblies. In this same study however, the authors concluded that the effect of allele phasing compared to the use of random alleles as with Hybpiper is not clear for phylogenetic reconstruction.

Assembled sequences for each sample were compiled, sorted by target sequence, and inspected for paralogs using the scripts ‘retrieve_sequences.py’, and ‘paralog_investigator.py’. Data on length of assembled coding sequences for each target sequence and statistics on assembly efficiency were calculated with the scripts ‘get_seq_lengths.py’ and ‘hybpiper_stats.py’. Coding sequences with paralog warnings were manually checked and potential paralogs were excluded from downstream analyses. Exons that are in the same loci were concatenated and we used Geneious v9.1.8. (Biomatters Ltd., Auckland, New Zealand) to create alignments for each loci with MAFFT v7.309 (Katoh & Standley, 2013), followed by manual inspection.

### Population structure across drainage basins

To examine population structure we ran ADMIXTURE v1.3 (Alexander *et al.*, 2009) and considered clustering scenarios with up to 6 groups (K=1–6). SNPs used as input were extracted from our target enrichment data with dDocent (Puritz *et al.*, 2014) by mapping the sequenced reads to the reference sequences of our targeted loci, assembling the reads, and calling SNPs. We further filtered our resulting SNPs to include loci with no missing data, only one SNP per locus, and a minor allele frequency of at least 0.05 using VCFtools (Danecek *et al.*, 2011). Given the high degree of missing data in the ddRADseq loci, we did not use these data for population structure inference.

### Phylogenomics and phylogenetic networks

#### Phylogenomics

A total of 42 samples were successful in the target enrichment experiment (four *Marathrum* samples failed sequencing or recovered relatively short exons of low quality. See Fig. S2), and these were used to conduct phylogenetic inference using concatenated and coalescent approaches. The concatenated target enrichment dataset and our *min17* ddRADseq assembly were analyzed under maximum likelihood (ML) with RAxML-HPC BlackBox v8.2.12 (Stamatakis, 2014) on the CIPRES Science Gateway. We ran 1000 bootstrap replicates to assess branch support and considered each locus as a separate partition in our target enrichment dataset. ddRADseq trees were rooted using the outgroup rooting of our target enrichment data as a reference. Gene tree inference was conducted using the TICR pipeline (Stenz *et al.*, 2015) to run MrBayes v3.2.7 (Ronquist & Huelsenbeck, 2003) for each target enrichment locus and generate 50% consensus majority rule trees. We specified the GTR+G model of substitution (Abadi *et al.*, 2019) with 3 runs, 4 chains, 2 million generations, a sample frequency of 200, and a burn-in fraction of 0.25. Species tree inference was conducted with ASTRAL III (Zhang *et al.*, 2018) and SVDQuartets (Chifman & Kubatko, 2014), both of which are coalescent-based approaches. Unrooted Bayesian gene trees were used as input for ASTRAL. We ran SVDQuartets analyses evaluating all quartets and using 1000 bootstraps on the concatenated sequences of our target capture data and the *min4* ddRADseq assembly. The latter comprises the maximum phylogenetic information necessary for a quartet based analysis (Eaton *et al.*, 2016) and is the data set that maximizes the number of shared loci recovered. The ddRADseq species trees were rooted using the outgroup rooting of our target enrichment data as a reference.

#### Phylogenetic networks

We explored non-bifurcating relationships in *Marathrum* by inferring phylogenetic networks in PhyloNet (Than *et al.*, 2008; Wen *et al.*, 2018). Following the results obtained with ADMIXTURE we randomly selected a subset of samples of *M. utile* and *M. foeniculaceum* from each drainage basin that did not show evidence of recent admixture. We kept those from Boyacá in order to infer how individuals in this population are related to Caribe and Antioquia. In addition, we conducted a second set of analyses that included admixed individuals expecting to find multiple reticulation events. For the latter, we created two datasets each of which included two individuals per population. We chose these individuals semi-randomly so that both “pure” and “admixed” individuals as inferred in ADMIXTURE were included. We rooted the previously inferred Bayesian consensus gene trees in R v4.0.2 with the outgroup *W. squamulosa* and inferred maximum pseudo-likelihood networks with 0–5 allowed reticulation events in PhyloNet. We followed Blair & Ané, 2020 in their recommendations for choosing among resulting phylogenetic networks by plotting the maximum number of allowed reticulations versus the resulting pseudolikelihood score, and selecting the number of reticulation events when a plateau begins to appear.

### Divergence times for populations across drainage basins

To provide the evolutionary context in which populations of *Marathrum* diverged across drainage basins, we inferred a time tree with BEAST 2.6.0 (Bouckaert *et al.*, 2019). We run two chains for 3 billion generations. A secondary calibration was defined for the stem node of *Marathrum*, using *Weddellina squamulosa* as an outgroup and a broad normal prior distribution with a mean of 62 Ma and a standard deviation of 4 Ma. This distribution encompassed the results for this node reported in previous published work (Magallón *et al.*, 2015; Ruhfel *et al.*, 2016) that utilized the fossil *Paleoclusia chevalieri* Crepet & Nixon for calibration. We assessed convergence of runs with Tracer v1.7.1 (Rambaut *et al.*, 2018), discarded a 10% burnin fraction, combined the log files and posterior distribution of trees from the two runs with LogCombiner, and generated a maximum clade credibility (MCC) tree with mean node heights with TreeAnnotator.

## Results

### Data processing: ddRADseq, genome assembly, and target enrichment

The number of loci shared across samples in our complete data set (n=95) and ipyrad assembly statistics for the subset of 34 samples (*min4* and *min17*) are in Tables S2 and S3. A total of 169,474 loci that are shared in minimum 4 individuals were recovered across all 95 *Marathrum* and *Weddellina* samples, but only 70 loci were recovered at 50% missing data) (Table S2). The *min4* (~88% missing data) and *min17* (50% missing data) assemblies of our subset of 34 samples resulted in 28,023 and 679 recovered loci respectively. Our results show that, although there are many loci recovered in our dataset, a smaller proportion of those are present across most samples, with no loci shared in all of them.

*De novo* assembly of genome-skimming reads resulted in a draft assembled genome of *M. utile* consisting of 627,733 contigs with a total length of 1.095 Gb and N50=11,454 bp (Bioproject PRJNA673497, SRR12956182). After running the pipeline Sondovac using the draft genome as a template for the identification of sequences to be targeted and applying our added filtering steps, we identified 536 sequences in 218 nuclear, single-copy loci. We recovered 208 loci across samples with an average of 507 exons in 205 loci. Figure S2 and Table S4 show the loci recovery efficiency and the number of sequences and loci recovered per sample respectively. Recovery of assembled sequences was lower for the outgroup taxa (*W. squamulosa*) but we included this sample for rooting purposes in downstream analyses.

Most warnings resulting from running paralog_investigator.py in Hybpiper corresponded to instances where identical contigs that differed only in length were assembled to a target sequence for a sample. In other instances, the differing contigs corresponded to alleles. In all such cases Hybpiper chooses the contig with the highest coverage. When contigs have similar sequencing depth, Hybpiper chooses one based on percent identity with the reference sequence (Johnson *et al.*, 2016). We excluded 28 exons where multiple contigs mapped to the same reference sequence, contained more than two alleles per SNP and included a high number of SNPs. By removing these exons, one locus was excluded.

### Population structure across drainage basins

Read assembly and SNP calling in dDocent recovered 203 variant sites. Parametric population structure estimation with ADMIXTURE resulted in a lower cross-validation error for *K*=4 and K=5 populations (0.08310 and 0.09195, respectively; Fig. **2b** and Fig. S3). The best model shows that within each Antioquia and Caribe there is strong population structure with two genetically distinct, non-admixed clusters as well as multiple individuals with admixture proportions from the two clusters (*i.e.* hybrids). In both drainage basins, the two distinct, non-admixed genetic clusters correspond to the phenotypes of *M. foeniculaceum* and *M. utile*. All admixed individuals have about equal proportions of ancestry of the non-admixed clusters. Furthermore, the admixed individuals have highly dissected leaves and their phenotype corresponds to that of *M. foeniculaceum* (Fig. **1b**). There is no admixture detected in individuals of the same species across drainage basins. The genetic constitution of the two individuals that represent the population from Boyacá was inferred to include relatively equal proportions from *M. utile* from Caribe and *M. foeniculaceum* from Antioquia. The only difference between K=4 and K=5 is that the latter detects population substructure in *M. utile* from Antioquia.

**Figure 2.**
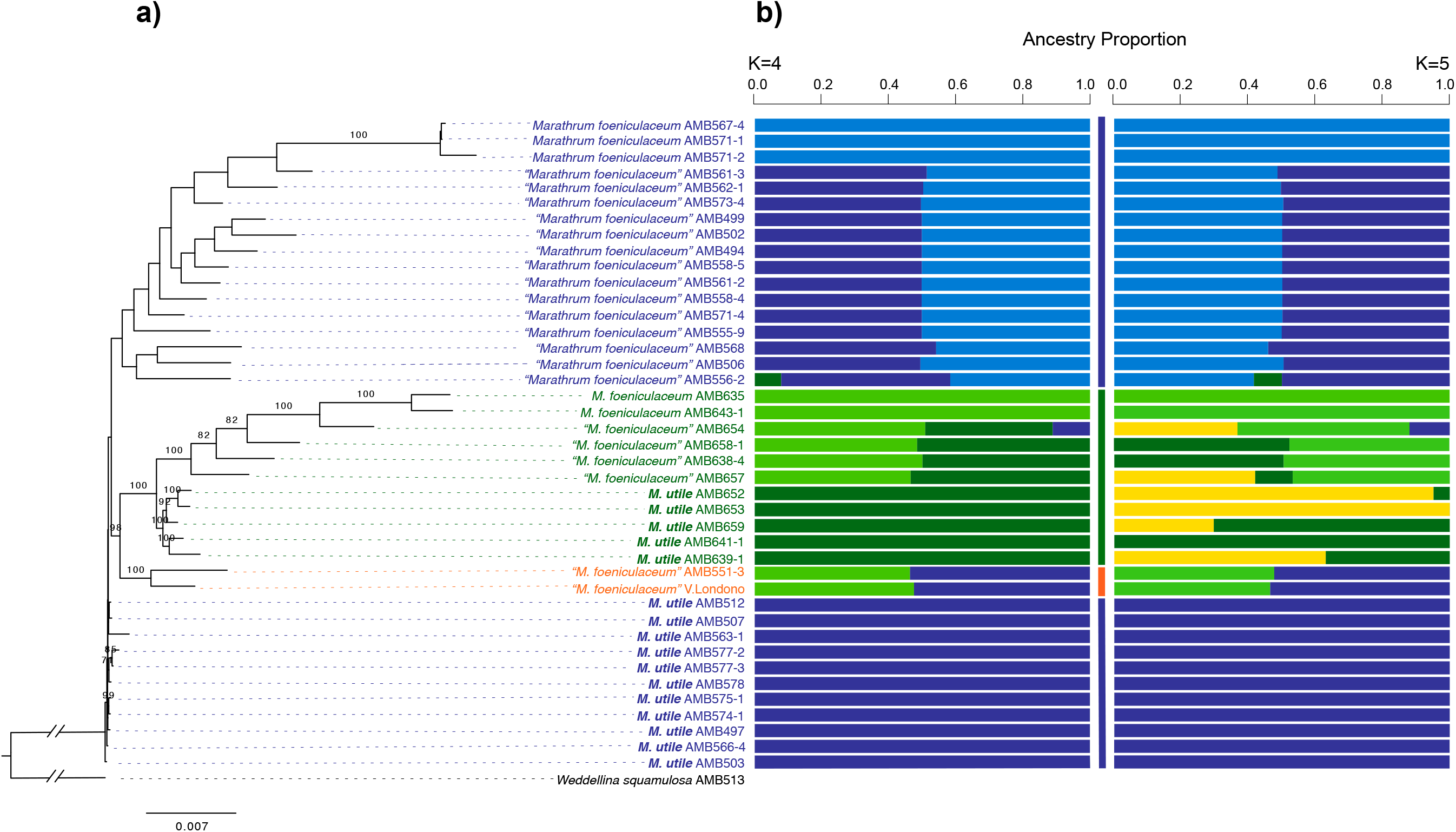
Phylogenetic relationships and ancestry proportion of individuals from Antioquia (green), Caribe (blue) and Boyacá (orange) inferred from target enrichment data. a) Maximum likelihood tree inferred from concatenated analyses of 207 targeted loci. Bootstrap support (≥70) is shown above branches. Admixed individuals are indicated with quotation marks. b) Ancestry proportions for K=4 and K=5 inferred with ADMIXTURE. Vertical bars indicate the drainage basin where each individual was collected.

### Phylogenomics and Phylogenetic networks

#### Phylogenomics

Results from ML analyses of all samples with target enrichment and ddRADseq data are shown in Figs. **2** and S4 respectively. Species trees for both datasets are shown in Fig. S5. Phylogenetic inference under the species coalescent with SVDQuartets and ASTRAL III was performed using individuals as tips (See Fig. S5) due to our prior evidence of the presence of admixed individuals **(Fig. 2b)**. The resulting trees do not recover species as clades, contradicting the current taxonomic treatment of *Marathrum*, and suggesting convergent evolution of forms in the genus. The target enrichment ML and ASTRAL III trees (Figs. **2** and S5b) are consistent, with Antioquia and Boyacá forming sister clades, but differ in that *M. utile* in Caribe is recovered as a clade in the species tree and as paraphyletic in the ML tree. Furthermore, target enrichment trees (Figs. **2** and S5) show *M. foeniculaceum* and *M. utile* within each driange basin as clades, except for the SVDQuartets tree (Fig. **2a**), where *M. foeniculaceum* from Antioquia is not monophyletic. The placement of Boyacá differs in this tree as well. Unlike the target enrichment trees, the trees inferred with ddRADseq data using ML (Fig. S4) and SVDQuartets (Fig. S5a) do not recover individuals with the phenotypes of *M. foeniculaceum* or *M. utile* in Caribe as clades. The ddRADseq trees differ in the placement of Boyacá and in the recovery of *M. foeniculaceum* from Antioquia as paraphyletic and monophyletic in the ML and SVDQuartets trees respectively. The latter tree exhibits weak support for most branches.

To explore and eliminate the conflict that can arise in phylogenetic reconstruction with the inclusion of admixed individuals that have ancestry proportions from *M. foeniculaceum* and *M. utile* inhabiting the same drainage basin, we removed all admixed individuals. However, we kept those from Boyacá in order to infer how individuals in this population are related to Caribe and Antioquia. We re-analyzed our data as described above, but we also conducted Bayesian inference (BI) in MrBayes v3.2.7 with 2 runs, 4 chains, 10 million generations, a sample frequency of 10,000, and a burn-in fraction of 0.25. Phylogenetic inference with ddRADseq on concatenated data was inferred from an ipyrad assembly with ~50% missing data, whereas species tree inference was conducted on an assembly with minimum four individuals per loci (Table S3 for ipyrad statistics). Each locus in our target enrichment dataset was set as a separate partition specifying the GTR+G substitution model for each partition (Abadi *et al.*, 2019). The GTR+G substitution model was also specified in the BI analysis of ddRADseq. We used the recovered genetic units from ADMIXTURE as the taxonomic units for species tree inference.

Results from these analyses are shown in Fig. **3**. These fundamentally contradict the results reported above using admixed and non-admixed individuals together. Maximum likelihood and BI trees of target enrichment data (Fig. **3a** and S6) and species trees for both datasets (Fig. **3b,c**) support the current taxonomic treatment of *Marathrum,* with *M. foeniculaceum* recovered as monophyletic and *M. utile* either as monophyletic (SVDQuartets inference in Fig. **3b**) or paraphyletic (ML, BI and ASTRAL III trees in Fig. **3a,c**, and S6). Plants from Boyacá, all of which have the phenotype of *M. foeniculaceum* (Fig. **1**), are found as sister to *M. foeniculaceum* from Antioquia and Caribe in all species tree inferences. Maximum likelihood (Fig. S6) and BI (Fig. **3a**) analyses of ddRADseq, which as mentioned before is characterized by high levels of missing data, show a biogeographic pattern where Antioquia evolves from Caribe and do not recover *M. foeniculaceum* and *M. utile* as clades. The reasons for the discordance among target enrichment and ddRADseq datasets are further discussed.

**Figure 3.**
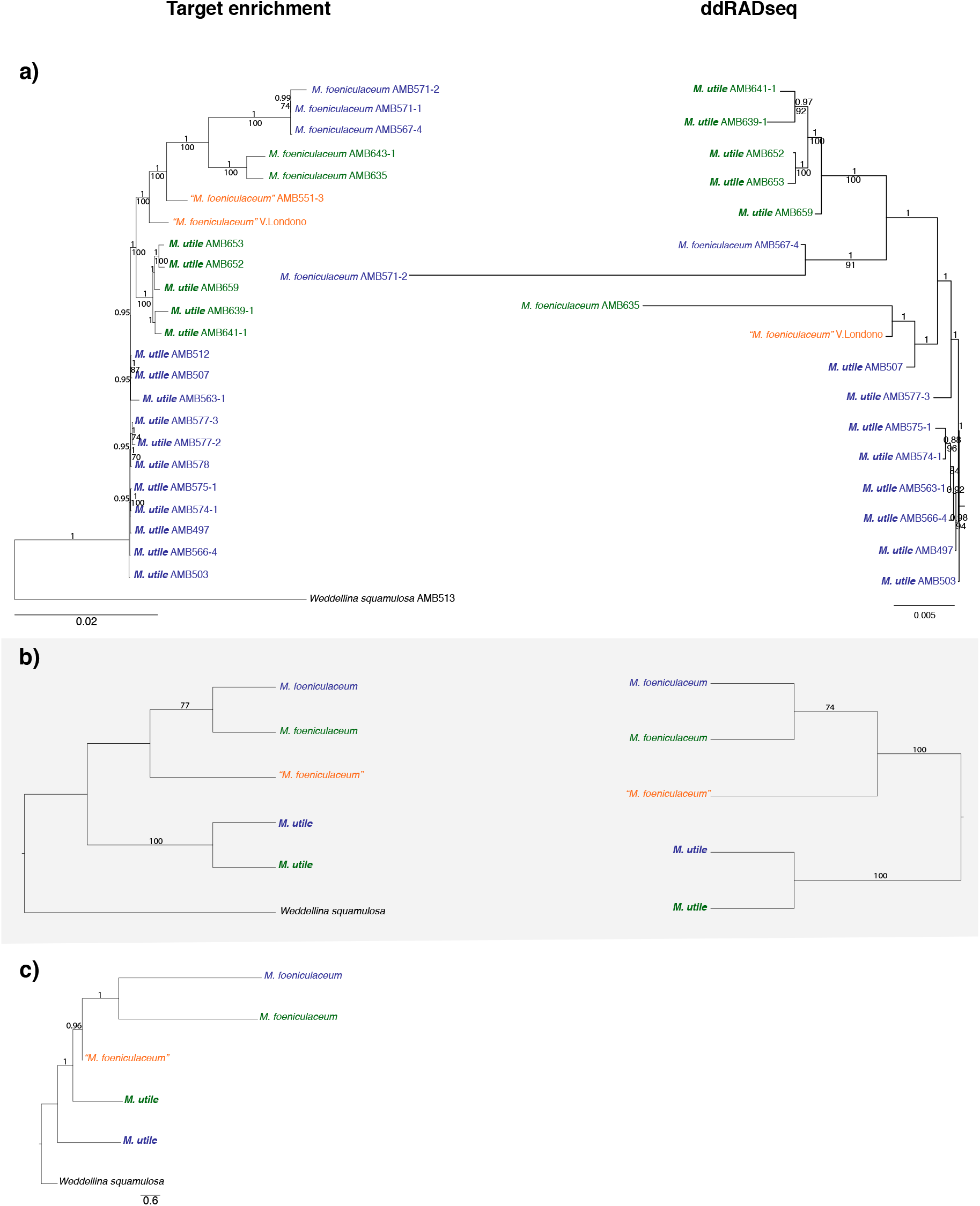
Phylogenetic trees inferred using data concatenation and species tree inference on target enrichment and ddRADseq data excluding admixed individuals from Antioquia and Caribe. The provenance of individuals is indicated in green (Antioquia), blue (Caribe), and orange (Boyacá). ddRADseq topologies are rooted based on results from target enrichment. Admixed individuals as inferred with ADMIXTURE are indicated with quotation marks. a) Majority rule consensus bayesian trees inferred from concatenated data. Posterior probabilities ≥0.7 are shown above branches. Bootstrap support (≥70) from ML reconstruction are shown below branches. b) Species tree inferred with SVDquartets. Quarter support values >70 shown above branches. c) Species tree inferred with ASTRAL III for the target enrichment dataset. Local posterior probabilities > 0.7 shown above branches.

#### Phylogenetic networks

Comparison of pseudolikelihood scores for 0–5 maximum reticulation events show that at two allowed reticulations, the change in pseudolikelihood begins to plateau (Fig. **4**). The phylogenetic network shows a reticulation event in the branch that leads to the two individuals from Boyacá, in line with our phylogenetic reconstructions excluding admixed individuals and our ADMIXTURE results. A second reticulation event is inferred deeper in the phylogeny with *M. foeniculaceum* as either sister to *M. utile* or originating from it. When including admixed individuals in our analysis, comparison of pseudolikelihood scores show that after 5 reticulation events, the change in pseudolikelihood has not plateaued (Fig. S7), suggesting that multiple admixture events have taken place in *Marathrum* in northern South America.

**Figure 4.**
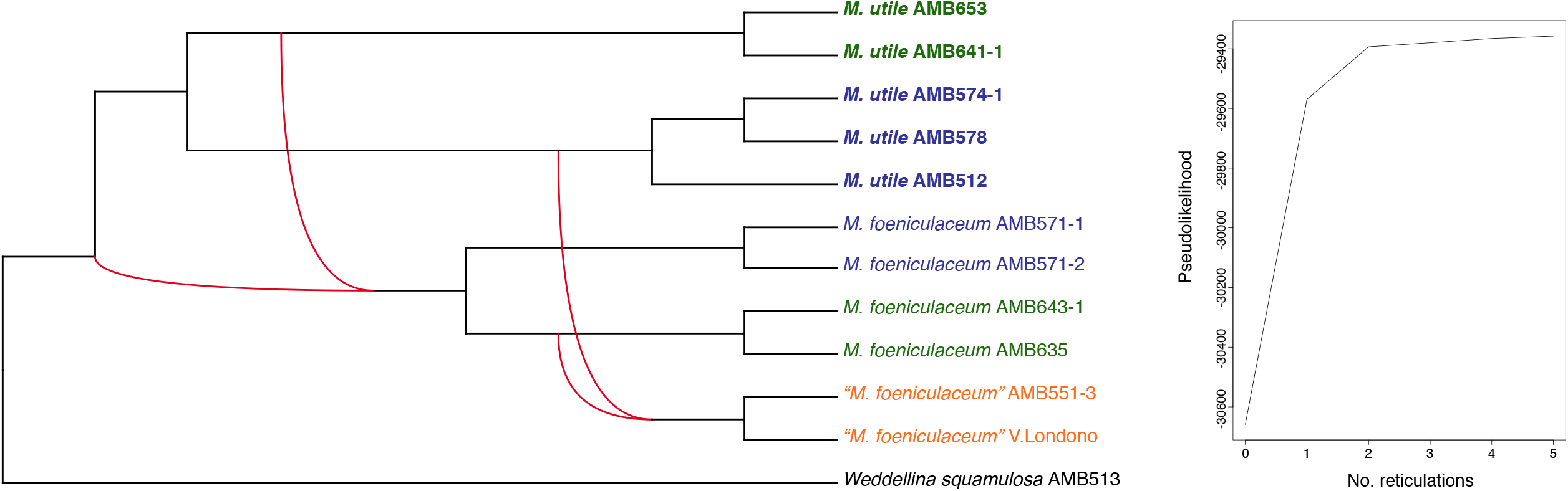
Phylogenetic network resulted from maximum pseudo-likelihood inference with PhyloNet. After 2 reticulation events, the pseudolikelihood score starts to plateau. Provenance of samples is shown in green (Antioquia), blue (Caribe), and orange (Boyacá). Admixed individuals as inferred with ADMIXTURE are indicated with quotation marks. Inferred reticulation events are depicted with red lines.

### Divergence times for populations across drainage basins

The BEAST calibrated phylogeny of *Marathrum* is shown in Fig. **5**. All populations sampled across the Andes and the SNSM are inferred to have shared a most recent common ancestor at ~4Ma at most. The topology supports the current taxonomic treatment of the group and is consistent with our phylogenetic trees obtained from target enrichment concatenated data and after excluding admixed individuals. Divergence times obtained in our study are compared to previous studies where the crown age of Podostemaceae was also estimated in Table 1.

**Figure 5.**
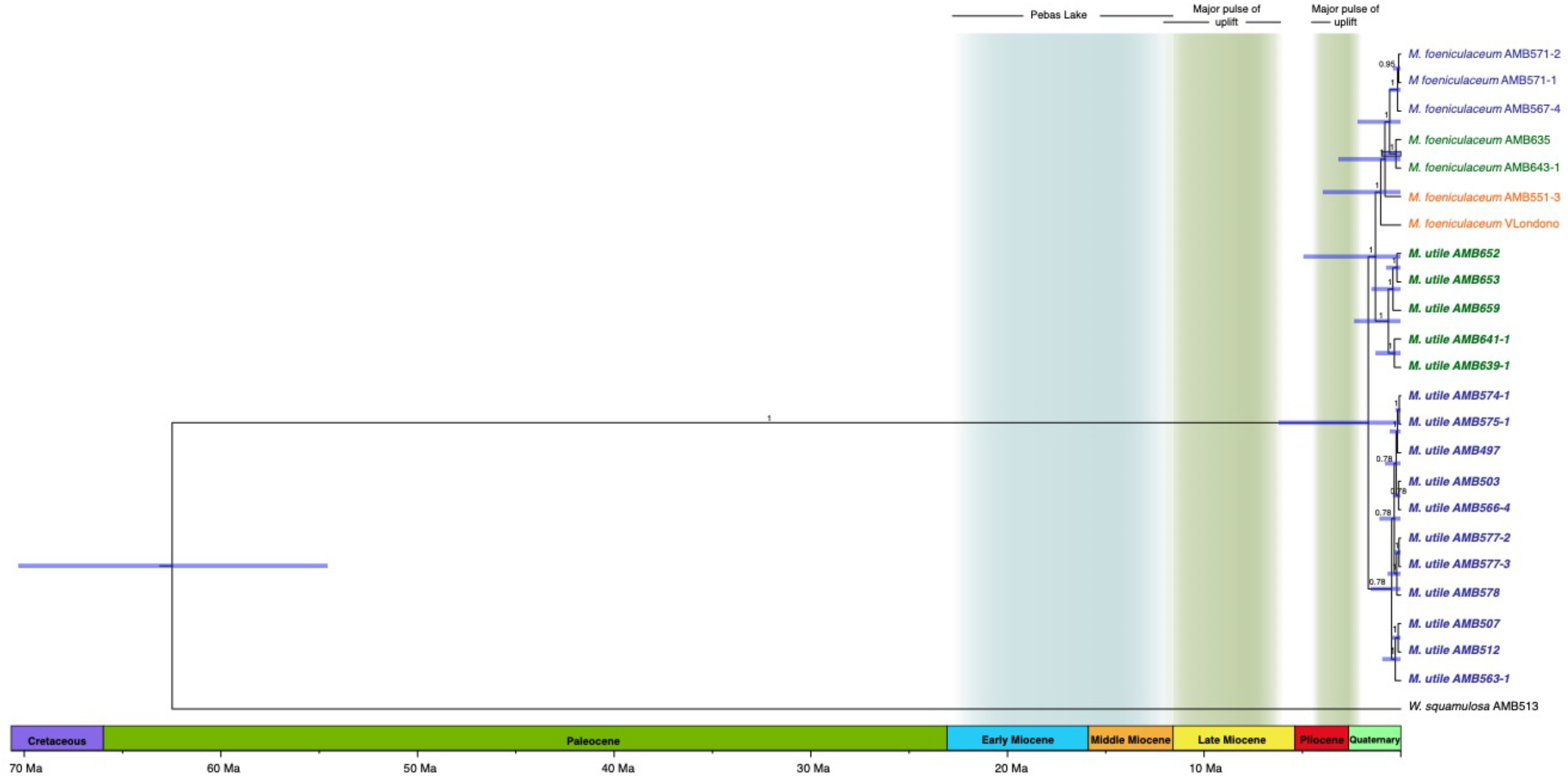
Divergence time tree inferred from *BEAST. Vertical bars indicate hypothesized landscape features and geological events. Blue horizontal bars indicate the 95% HPD. Posterior probabilities are shown above branches. The provenance of individuals is indicated in green (Antioquia), blue (Caribe), and orange (Boyacá).

**Table 1.** Results of our calibration and the calibrations for the same node inferred in two previous studies. Ruhfel et al., presented two dating results for alternative placements of the fossil *Paleoclusia chevalieri*

|  | Bedoya et al., 2020 | Magallón et al., 2015 | Ruhfel et al., 2016 |  |
| --- | --- | --- | --- | --- |
| 95% HPD Min (Ma) | 54.56 | 53.53 | 49.9 | 55.4 |
| 95% HPD Mean (Mya) | 62.43 | 67.82 | 62 | 66.1 |
| 95% HPD Max (Mya) | 70.30 | 77.29 | 71.9 | 76.8 |

## Discussion

Together, our population structure, phylogenomic, phylogenetic networks, and divergence-dating analyses inform us about how through the Miocene and Pliocene, landscape change in northern South America shaped the evolution of *Marathrum* and of river basins. We show that during this time period, populations of both *M. foeniculaceum* and *M. utile* split into different drainage basins currently interrupted by the Andes, the SNSM, and inter-Andean valleys and became genetically distinct (Figs. **3**,**5**). As a result, strong population structure consistent with geography was inferred in populations of each species (Fig. **2**). At the same time we demonstrate that there is little to no gene flow in populations of the same species across drainage basins (Fig. **2**) and therefore, that the Andean cordilleras, the SNSM, and/or inter-Andean valleys currently act as barriers that limit gene flow in this particular group of plants. We found that while populations across drainage basins became genetically distinct, the sympatric distribution of *M. foeniculaceum* and *M. utile* resulted in hybridization within drainage basins, and we report that hybrids show the phenotype of only *M. foeniculaceum*. The current taxonomic treatment of the genus is validated in this study. Divergence dating results imply that fast-flowing water ecosystems in Antioquia, Caribe, and Boyacá and populations of *Marathrum* inhabiting them became isolated (Fig. **5**) very recently in geologic time (< 4Ma). We propose that the phylogenetic relationships of populations (namely Boyacá as sister to Caribe-Antioquia) reflect the scenario for the divergence of drainage basins as the Andes uplifted (Fig. **3**).

### Population structure across drainage basins

#### Barriers to gene flow

In addition to mountains and valleys being a barrier to gene flow in populations of *M. foeniculaceum and M. utile* in northern South America, a limited dispersal ability of pollen or seeds in these plants could contribute to the strong population structure and lack of within species gene flow reported here across drainage basins (Fig. **2**). This is in line with previous work on Podostemaceae, where high levels of differentiation among populations of *Hydrobium japonicum* and *Cladopus doianus* in Japan and of *Podostemum irgangii* in Brazil were attributed to be the result of reduced dispersal due to their restricted habitat (Baggio *et al.*, 2013; Katayama *et al.*, 2016). This pattern is also seen in freshwater fishes (which share the same habitat as *Marathrum*) where range evolution is generally constrained by watershed boundaries (Albert & Crampton, 2010; Lujan & Armbruster, 2011; Musilová *et al.*, 2013; Picq *et al.*, 2014; Roxo *et al.*, 2014; Tagliacollo *et al.*, 2015; Lujan *et al.*, 2019; Albert *et al.*, 2020).

However, long-distance dispersal has been invoked as the explanation for the pantropical distribution of Podostemaceae (Kita & Kato, 2004; Koi *et al.*, 2015; Ruhfel *et al.*, 2016). Birds may be possible vectors for seed dispersal in Podostemaceae if seeds get attached to their feet via a sticky mucilage in the seed coat (Willis, 1902; Philbrick & Novelo, 1995, 1997; Leleeka *et al.*, 2016). Future work should focus on the extent of dispersal of Podostemaceae within a single river and across rivers to determine how seed and pollen dispersal is facilitated or hindered.

#### Genetic drift

Populations of *Marathrum* and of Podostemaceae in general tend to be small, with a discontinuous distribution due to their high degree of habitat specificity (Willis, 1914; van Royen, 1951; Rutishauser, 1995; Cook, 1996; Philbrick *et al.*, 2010; Koi *et al.*, 2015; Cheek & Lebbie, 2018). We believe that in such small populations with short generation times, genetic drift may be a rather strong evolutionary force. As drainage basins were established, this mechanism would have acted independently in populations of each species within each drainage basin, quickly fixing their alleles, and resulting in homogeneous populations that are genetically distinct across drainage basins as found in this study for *Marathrum* (Fig. **2**).

#### Hybridization scenarios and phenotype

Luna *et al.*, 2012 mentioned hybridization as a possible explanation for the existence of intermediate morphotypes among closely related species of *Marathrum* that are morphologically similar. In that same study however, the authors concluded that cross-compatibility is between morphological variants of the same species which had been previously synonymized under *M. foeniculaceum* (Novelo *et al.*, 2009; Tippery *et al.*, 2011). Our study includes key insights into the reproductive biology of *Marathrum* by providing clear genomic evidence for hybridization without introgression between two phenotypically and phylogenetically distinct species in *Marathrum.*

We propose two hypotheses for the uniformity in admixture proportions across admixed individuals in Antioquia and Caribe (Fig. **2**). The genetic constitution of admixed individuals in these two drainage basins stems from 1) very recent and frequent admixture events (*i.e*. F1 hybrids of the two species with no further mixing or backcrossing) or 2) a past hybridization event followed by genome duplication resulting in allopolyploids. Under the latter scenario, hybridization needs to have occurred at minimum once in each river basin but not every generation. Fertile allopolyploids would reproduce the admixed genomes (and the *M. foeniculaceum* morphotype). Variation in ploidy and cell DNA content are unknown for these populations.

The dual ancestry of the two individuals that represent the population from Boyacá stems from *M. utile* from Caribe and *M. foeniculaceum* from Antioquia in more or less equal proportions (Fig. **2**). No individuals of *M. utile* and *M. foeniculaceum* are known to exist in or close to that population. Sampling needs to be expanded to confirm this pattern in more individuals in Boyacá, and to test whether admixture in individuals of this population is a result of recent hybridization with or without long distance transport of pollen or seeds, retention of ancestral polymorphism in this population, or allopolyploidy.

This study spotlights an interesting case of inheritance of a phenotypic trait: leaf shape. Given that all admixed individuals and putative hybrids show the phenotype of *M. foeniculaceum* (Figs. **1** and **2**), a likely hypothesis is that leaf shape is controlled by one or few genes, with highly dissected leaves being the dominant trait. Common garden experiments could test the hybrid origin of the admixed individuals and the genetic basis of leaf morphology.

### Phylogenomics

#### Incongruence in ddRAdseq vs target enrichment datasets with concatenated data

Missing data, internal short branches like those shown in our inferred trees in Figs. **2**, **3a**, and S4 and bioinformatic steps used for the assembly of loci (e.g. clustering threshold), have a strong impact on tree inference (Leaché *et al.*, 2015). These three phenomena are characteristic of our ddRADseq data and may be responsible for the discordance among datasets in this study (Figs. **3a** S5, and S6). As shown in Table S3, our ddRADseq data are characterized by high levels of missing data, possibly as a result of previously suggested high rates of evolution in Podostemaceae (Bedoya *et al.*, 2019). Rapid evolution would result in high sequence divergence, and allelic dropout would reduce the number of shared loci recovered (Wagner *et al.*, 2013; Arnold *et al.*, 2013; Eaton *et al.*, 2016). Incomplete lineage sorting (ILS) could be responsible for the recovery of the two individuals from Boyacá as paraphyletic in the tree inferred from the concatenated data using target enrichment data (Fig. **3a**), but more individuals should be sampled from this population to support this claim. When ILS is accounted for with quartet analysis of our *min4* ddRADseq data, phylogenetic inference is congruent with the trees inferred from target enrichment. The latter trees are free from the biases mentioned above and therefore provide more reliable results for discussing the evolutionary history of *Marathrum.*

#### Systematics of Marathrum

Our results demonstrate that *M. foeniculaceum* and *M. utile* are consistent with traditional taxonomic units (Fig. **3**), being most likely both monophyletic as inferred with SVDQuartets, or an ancestor-descendant species pair as resulting from ASTRAL III analyses. Even though both ASTRAL III and SVDQuartets consider ILS as a source for gene tree incongruence, species tree inference may be inconsistent when gene trees are estimated from data for loci of finite length (Wascher & Kubatko, 2020) as is the case in ASTRAL III. In contrast, SVDQuartets has been confirmed to be statistically consistent under various scenarios of ILS (Wascher & Kubatko, 2020). Our phylogenetic reconstructions are affected by the inclusion of admixed individuals (Figs. **2**, **3,** and S4). This is because hybrids contain alleles originating from both parents (Bastide *et al.*, 2018; Zhang *et al.*, 2019) causing the parental lineages and hybrids to cluster together. This phenomenon has been reported across various groups (McVay *et al.*, 2017; García *et al.*, 2017; Zhang *et al.*, 2019; Ginsberg *et al.*, 2019) and affects both concatenated and species tree inference (Leaché *et al.*, 2014).

### Landscape change, river basin formation and timing of divergence of populations of *Marathrum* in nSA

With the evidence presented above, we propose that the pattern of divergence of Antioquia, Boyacá, and Caribe tracks the scenario in which drainage basins split from one another. Based on our species tree inferences, Boyacá (eastern slope of the EC) diverged earlier from Caribe and Antioquia. Even though the SVDQuartets species tree shows weak support for the placement of *M. foeniculaceum* from Boyacá, both this and the ASTRAL III (Fig. **3b,c**) show the same biogeographic pattern. Reticulation events (Fig. **4**) could have taken place in sympatry or be the result of vicariance followed by secondary contact mediated by, for example, river capture events or less likely, long distance dispersal.

Several studies document that major pulses of uplift took place during the Late Miocene and Pliocene asynchronously across the three Andean cordilleras at *ca.* 12–6 Ma and *ca.* 4.5 Ma, resulting in the current elevation of the Andes (Gregory-Woodzicki, 2000; Garzione *et al.*, 2008; Mora *et al.*, 2010; Hoorn *et al.*, 2010). These dates are contested by geological studies of mainly coarse-grained deposits that suggest that the Eastern Cordillera was already well underway in Eocene times and nearly complete in the Late Miocene (>11 Ma) (Cooper *et al.*, 1995; Montes *et al.*, 2019; Rodríguez-Muñoz *et al.*, 2020). Our divergence dating analysis results in Fig. **5** imply that the orogenic history of the Andean Cordillera and inter-Andean valley formation proposed by either of the two hypotheses, did not result in the complete separation of the currently isolated, genetically distinct populations of *M. foeniculaceum* and *M. utile* in Antioquia, Boyacá and Caribe until < 5 Ma.

Furthermore, we favor the hypothesis of rapid pulses of Andean uplift in the Late Miocene and Pliocene by showing that the establishment of completely isolated fast-flowing water ecosystems in Antioquia, Caribe and Boyacá was not completed until < 4Ma (Fig. **5**) when the Andes, SNSM, and inter-Andean valleys became a barrier to gene flow. Our results are in line with Anderson *et al.*, 2016 who using U-Pb detrital zircon provenance, petrographic and paleoprecipitation data, conclude that the northern Andes was not a substantial orographic river barrier until *ca.* 3–6 Ma when rapid uplift took place. A lowland connection between the Amazon, Caribe and Magdalena drainage basins is corroborated by the affinity of fossils in Miocene and Pliocene deposits found across drainage basins (Lundberg & Chernoff, 1992; Aguilera *et al.*, 2013). The short branches and very recent splits in populations in Caribe indicate that the rivers in this drainage basin originated and split very recently in geologic time (Figs. **3**, **5**, and S4). Indeed, surface uplift that gave rise to the current topology of the SNSM has been proposed to have taken place no earlier than 2 Ma (Villagómez *et al.*, 2011; Parra *et al.*, 2019).

Since our calibrated phylogeny was inferred from concatenated data, the dates inferred for each of the nodes could be more recent than if species tree inference was used instead. However, the joint inference of gene trees and a calibrated species tree in *BEAST is computationally intensive and feasible only for moderate-size datasets (McCormack *et al.*, 2013). Divide-and-conquer methods must be used to improve scalability in species tree estimation in *BEAST (Zimmermann *et al.*, 2014) if hundreds of loci are to be considered, potentially increasing the uncertainty in the estimation of node ages. This study represents a significant advance in our understanding of the impact of landscape change in the evolution of plants in rivers in northern South America and provides a comprehensive genomic dataset for the only plant genus that inhabits fast-flowing water ecosystems across the Western, Central, and Eastern Andean cordilleras and the Sierra Nevada de Santa Marta.

## Acknowledgements

We are thankful to Dr. C.Thomas Philbrick for his helpful and thorough insights into the biology and classification of Podostemaceae, access to voucher specimens, and help with determination of samples. Special thanks to Dr. Santiago Madriñán at the Universidad de los Andes and the Jardín Botánico de Cartagena “Guillermo Piñeres”, Turbaco, Colombia for facilitating field work logistics and providing facilities for specimen processing in Colombia. We thank Dr. Camilo Montes at the Universidad del Norte, Barranquilla, Colombia for insightful conversations on the geological background of nSA, and Dr. Camila Martínez at the Smithsonian Institution in Panama for comments on the manuscript. Maria Paula Contreras, David Ocampo, Daniel Ocampo, Jules Dominé, Fernando Bedoya, and Adrián Pinzón provided invaluable assistance in the field. Viviana Londoño provided a sample of *M. foeniculaceum*. Voucher specimens are deposited at the UNIANDES herbarium of the Universidad de los Andes, Bogota, Colombia. Samples were collected under the MARCO collecting permit Res. No. 1177 October 9, 2014 of the Universidad de los Andes. Funding was provided by the Botanical Society of America and American Society of Plant Taxonomists Research Awards, Explorers Club Exploration Fund Grant, Department of Biology at the University of Washington Giles Award and WRF Hall International Endowed Fellowship, UW Herbarium Endowment, UW Graduate School Boeing International Fellowship, and the Colciencias fellowship for Graduate studies (Doctorados en el Exterior-679) to A.M.B. This work used the Vincent J. Coates Genomics Sequencing Laboratory at UC Berkeley, supported by NIH S10 OD018174 Instrumentation Grant.

## Author Contribution

A.M.B. and R.G.O designed the study. A.M.B. collected and analyzed the data; R.G.O. and A.D.L. provided guidance on the methods for data analyses and interpretation of results, as well as reagents, equipment, and laboratory space. A.M.B. wrote the manuscript with significant contributions from R.G.O and A.D.L.

## Data Availability

All raw sequence reads from the low-coverage genome, target enrichment experiment, and ddRADseq datasets are deposited in the National Center for Biotechnology Information (NCBI) Sequence Read Archive under Bioproject PRJNA673497 (available upon initial acceptance). Fasta sequences of targeted loci, processed data and all trees generated in this study will be deposited in a public repository upon initial acceptance.

## Supplementary Materials

**Table S1.** Voucher numbers, NCBI accessions and locations for the samples collected and included in this study. Samples are arranged by species. Admixed individuals are indicated in quotations marks, and individuals that have the phenotype of *M. foeniculaceum* but whose genetic constitution was not assessed with admixture are indicated as morphs since they could correspond to *M. foeniculaceum* or to hybrids. The sample used for generating genome-skimming data is in bold (SRR12956182 for WGS read data). Genomic data are deposited in the NCBI Sequence Read Archive under Bioproject PRJNA673497.

| Collector No | SRA accession #<br>ddRADseq | SRA accession #<br>target enrichment | GenBank accession<br>numbers | Species | Coordinates | Basin/River |
| --- | --- | --- | --- | --- | --- | --- |
| **AMB 497** | **SRR12971533** | **SRR12975804** | - | *Marathrum utile* | **11°11'47.07"N, 73°27'35.53"W** | **Caribe/Ancho** |
| AMB 503 | SRR12989095 | SRR12983242 | - | *Marathrum utile* | 11°11'33.46"N, 73°34'59.01"W | Caribe/Chorro Catalina |
| AMB 507 | SRR12971532 | SRR12975803 | - | *Marathrum utile* | 11°6'13.93"N, 73°52'19.28"W | Caribe/Buritaca |
| AMB 508 | SRR12971530 | - | - | *Marathrum utile* | 11°6'13.93"N, 73°52'19.28"W | Caribe/Buritaca |
| AMB 509 | SRR12971529 | - | - | *Marathrum utile* | 11°6'13.93"N, 73°52'19.28"W | Caribe/Buritaca |
| AMB 512 | FAILED | SRR12975802 | - | *Marathrum utile* | 11°5'34.16"N, 73°53'12.51"W | Caribe/Buritaca |
| AMB 563-1 | SRR12971527 | SRR12975801 | - | *Marathrum utile* | 11°13'30"N, 73°41'50"W | Caribe/Don Diego |
| AMB 563-3 | SRR12971526 | - | - | *Marathrum utile* | 11°13'30"N, 73°41'50"W | Caribe/Don Diego |
| AMB 563-4 | SRR12971525 | - | - | *Marathrum utile* | 11°13'30"N, 73°41'50"W | Caribe/Don Diego |
| AMB 566-1 | SRR12971524 | - | - | *Marathrum utile* | 11°12'2"N, 73°27'42"W | Caribe/Ancho |
| AMB 566-3 | SRR12971523 | - | - | *Marathrum utile* | 11°12'2"N, 73°27'42"W | Caribe/Ancho |
| AMB 566-4 | SRR12971522 | SRR12975800 | - | *Marathrum utile* | 11°12'2"N, 73°27'42"W | Caribe/Ancho |
| AMB 566-6 | SRR12971521 | - | - | *Marathrum utile* | 11°12'2"N, 73°27'42"W | Caribe/Ancho |
| AMB 566-8 | SRR12971519 | - | - | *Marathrum utile* | 11°12'2"N, 73°27'42"W | Caribe/Ancho |
| AMB 566-9 | SRR12971518 | - | - | *Marathrum utile* | 11°12'2"N, 73°27'42"W | Caribe/Ancho |
| AMB 566-14 | SRR12971517 | - | - | *Marathrum utile* | 11°12'2"N, 73°27'42"W | Caribe/Ancho |
| AMB 566-18 | SRR12971516 | - | - | *Marathrum utile* | 11°12'2"N, 73°27'42"W | Caribe/Ancho |
| AMB 574-1 | SRR12971515 | SRR12975799 | - | *Marathrum utile* | 11°12'39"N, 73°15'8"W | Caribe/Jerez |
| AMB 574-2 | SRR12971514 | - | - | *Marathrum utile* | 11°12'39"N, 73°15'8"W | Caribe/Jerez |
| AMB 575-1 | SRR12971513 | SRR12975798 | - | *Marathrum utile* | 11°12'39"N, 73°15'8"W | Caribe/Jerez |
| AMB 575-2 | SRR12971512 | - | - | *Marathrum utile* | 11°12'39"N, 73°15'8"W | Caribe/Jerez |
| AMB 575-3 | SRR12971511 | - | - | *Marathrum utile* | 11°12'39"N, 73°15'8"W | Caribe/Jerez |
| AMB 577-1 | SRR12971510 | - | - | *Marathrum utile* | 10°42'25"N, 73°17'4"W | Caribe/Jerez |
| AMB 577-2 | FAILED | SRR12975797 | - | *Marathrum utile* | 10°42'25"N, 73°17'4"W | Caribe/Badillo |
| AMB 577-3 | SRR12971507 | SRR12975796 | - | *Marathrum utile* | 10°42'25"N, 73°17'4"W | Caribe/Badillo |
| AMB 578 | - | SRR12983243 | - | *Marathrum utile* | 10°30'5"N, 73°16'9"W | Caribe/Guatapuri |
| AMB 639-1 | SRR12971506 | SRR12975795 | - | *Marathrum utile* | 6°16'0"N, 75°1'45"W | Antioquia/Arenal |
| AMB 639-2 | SRR12971505 | - | - | *Marathrum utile* | 6°16'0"N, 75°1'45"W | Antioquia/Arenal |
| AMB 639-3 | SRR12971504 | - | - | *Marathrum utile* | 6°16'0"N, 75°1'45"W | Antioquia/Arenal |
| AMB 639-4 | SRR12971503 | - | - | *Marathrum utile* | 6°16'0"N, 75°1'45"W | Antioquia/Arenal |
| AMB 639-5 | SRR12971502 | - | - | *Marathrum utile* | 6°16'0"N, 75°1'45"W | Antioquia/Arenal |
| AMB 641-1 | SRR12971501 | SRR12975793 | - | *Marathrum utile* | 6°13'57"N, 74°54'58"W | Antioquia/Arenal |
| AMB 641-2 | SRR12971500 | - | - | *Marathrum utile* | 6°13'57"N, 74°54'58"W | Antioquia/Arenal |
| AMB 641-3 | SRR12971499 | - | - | *Marathrum utile* | 6°13'57"N, 74°54'58"W | Antioquia/Arenal |
| AMB 642-1 | FAILED | FAILED | - | *Marathrum utile* | 6°13'57"N, 74°54'58"W | Antioquia/Arenal |
| AMB 642-2 | SRR12971497 | - | - | *Marathrum utile* | 6°13'57"N, 74°54'58"W | Antioquia/Arenal |
| AMB 651 | SRR12971596 | - | - | *Marathrum utile* | 5°55'58"N, 75°5'45"W | Antioquia/Santo Domingo |
| AMB 652 | SRR12971595 | SRR12975792 | - | *Marathrum utile* | 5°55'58"N, 75°5'45"W | Antioquia/Santo Domingo |
| AMB 653 | SRR12971594 | SRR12975791 | - | *Marathrum utile* | 5°54'51"N, 75°4'28"W | Antioquia/Santo Domingo |
| AMB 659 | SRR12971593 | SRR12975790 | - | *Marathrum utile* | 6°0'2"N, 74°56'15"W | Antioquia/Samaná Norte |
| AMB 567-4 | SRR12971578 | SRR12975808 | - | *Marathrum foeniculaceum* | 11°12'2"N, 73°27'42"W | Caribe/Ancho |
| AMB 571-1 | FAILED | SRR12975807 | - | *Marathrum foeniculaceum* | 11°13'46"N, 73°34'40"W | Caribe/Palomino |
| AMB 571-2 | SRR12971576 | SRR12975806 | - | *Marathrum foeniculaceum* | 11°13'46"N, 73°34'40"W | Caribe/Palomino |
| AMB 635 | SRR13013619 | SRR12983247 | - | *Marathrum foeniculaceum* | 6°16'4"N, 75°1'49"W | Antioquia/Arenal |
| AMB 643-1 | - | SRR12983246 | - | *Marathrum foeniculaceum* | 6°13'57"N, 74°54'58"W | Antioquia/Arenal |
| AMB 494 | SRR12971587 | SRR12975817 | - | "*Marathrum foeniculaceum*" | 11°13'7.63"N, 73°42'0.14"W | Caribe/Don Diego |
| AMB 499 | SRR12971586 | SRR12975816 | - | "*Marathrum foeniculaceum*" | 11°12'22.95"N, 73°15'17.26"W | Caribe/Jerez |
| AMB 502 | SRR12971575 | SRR12975805 | - | "*Marathrum foeniculaceum*" | 11°11'33.46"N, 73°34'59.01"W | Caribe/Chorro Catalina |
| AMB 506 | SRR13013618 | SRR12983248 | - | "*Marathrum foeniculaceum*" | 11°16'33.96"N, 73°55'26.57"W | Caribe/ Piedras |
| AMB 551-3 | - | SRR12983244 | - | "*Marathrum foeniculaceum*" | 4°53'29"N, 73°16'31"W | Boyacá/Batá |
| AMB 555-9 | SRR12971564 | SRR12975794 | - | "*Marathrum foeniculaceum*" | 11°16'33.96"N, 73°55'26.57"W | Caribe/Piedras |
| AMB 556-2 | SRR12971553 | SRR12975789 | - | "*Marathrum foeniculaceum*" | 11°16'0"N, 73°51'56"W | Caribe/MendiHuaca |
| AMB 558-4 | SRR12971542 | SRR12975788 | - | "*Marathrum foeniculaceum*" | 11°14'41"N, 73°50'35"W | Caribe/Guachaca |
| AMB 558-5 | SRR12971531 | SRR12975787 | - | "*Marathrum foeniculaceum*" | 11°14'41"N, 73°50'35"W | Caribe/Guachaca |
| AMB 561-2 | SRR12971520 | SRR12975786 | - | "*Marathrum foeniculaceum*" | 11°14'11"N, 73°41'51"W | Caribe/Don Diego |
| AMB 561-3 | SRR12971509 | SRR12975785 | - | "*Marathrum foeniculaceum*" | 11°14'11"N, 73°41'51"W | Caribe/Don Diego |
| AMB 562-1 | SRR12971498 | SRR12975784 | - | "*Marathrum foeniculaceum*" | 11°13'56"N, 73°42'2"W | Caribe/Don Diego |
| AMB 568 | SRR12971585 | SRR12975815 | - | "*Marathrum foeniculaceum*" | 11°12'2"N, 73°27'42"W | Caribe/Ancho |
| AMB 571-4 | SRR12971584 | SRR12975814 | - | "*Marathrum foeniculaceum*" | 11°13'46"N, 73°34'40"W | Caribe/Palomino |
| AMB 573-4 | FAILED | SRR12975813 | - | "*Marathrum foeniculaceum*" | 11°12'39"N, 73°15'8"W | Caribe/Jerez |
| AMB 638-4 | SRR12971582 | SRR12975812 | - | "*Marathrum foeniculaceum*" | 6°16'0"N, 75°1'45"W | Antioquia/Arenal |
| AMB 654 | SRR12971581 | SRR12975811 | - | "*Marathrum foeniculaceum*" | 5°54'51"N, 75°4'28"W | Antioquia/Santo Domingo |
| AMB 657 | SRR13013620 | SRR12983245 | - | "*Marathrum foeniculaceum*" | 5°54'56"N, 74°58'45"W | Antioquia/Samaná Norte |
| AMB 658-1 | SRR12971580 | SRR12975810 | - | "*Marathrum foeniculaceum*" | 6°0'2"N, 74°56'15"W | Antioquia/Samaná Norte |
| V.Londono | SRR12971579 | SRR12975809 | - | "*Marathrum foeniculaceum*" | 4°53'29"N, 73°16'31"W | Boyacá/Batá |
| AMB 492 | FAILED | - | - | *Marathrum foeniculaceum* (morph) | 11°13'7.63"N, 73°42'0.14"W | Caribe/Don Diego |
| AMB 493 | SRR12971574 | - | - | *Marathrum foeniculaceum* (morph) | 11°13'7.63"N, 73°42'0.14"W | Caribe/Don Diego |
| AMB 495 | FAILED | - | - | *Marathrum foeniculaceum* (morph) | 11°13'19.31"N, 73°31'38.01"W | Caribe/San Salvador |
| AMB 496 | SRR12971573 | - | - | *Marathrum foeniculaceum* (morph) | 11°13'19.31"N, 73°31'38.01"W | Caribe/San Salvador |
| AMB 498 | FAILED | - | - | *Marathrum foeniculaceum* (morph) | 11°11'47.07"N, 73°27'35.53"W | Caribe/Ancho |
| AMB 500 | SRR12971572 | - | - | *Marathrum foeniculaceum* (morph) | 11°12'22.95"N, 73°15'17.26"W | Caribe/Jerez |
| AMB 551-4 | SRR12971571 | - | - | *Marathrum foeniculaceum* (morph) | 4°53'29"N, 73°16'31"W | Boyacá/Batá |
| AMB 551-5 | SRR12971570 | - | - | *Marathrum foeniculaceum* (morph) | 4°53'29"N, 73°16'31"W | Boyacá/Batá |
| AMB 551-6 | SRR12971569 | - | - | *Marathrum foeniculaceum* (morph) | 4°53'29"N, 73°16'31"W | Boyacá/Batá |
| AMB 551-7 | SRR12971568 | - | - | *Marathrum foeniculaceum* (morph) | 4°53'29"N, 73°16'31"W | Boyacá/Batá |
| AMB 551-8 | SRR12971567 | - | - | *Marathrum foeniculaceum* (morph) | 4°53'29"N, 73°16'31"W | Boyacá/Batá |
| AMB 551-10 | SRR12971566 | - | - | *Marathrum foeniculaceum* (morph) | 4°53'29"N, 73°16'31"W | Boyacá/Batá |
| AMB 556-1 | SRR12971565 | - | - | *Marathrum foeniculaceum* (morph) | 11°16'0"N, 73°51'56"W | Caribe/MendiHuaca |
| AMB 556-3 | SRR12971563 | - | - | *Marathrum foeniculaceum* (morph) | 11°16'0"N, 73°51'56"W | Caribe/MendiHuaca |
| AMB 556-4 | SRR12971562 | - | - | *Marathrum foeniculaceum* (morph) | 11°16'0"N, 73°51'56"W | Caribe/MendiHuaca |
| AMB 556-5 | SRR12971561 | - | - | *Marathrum foeniculaceum* (morph) | 11°16'0"N, 73°51'56"W | Caribe/MendiHuaca |
| AMB 558-1 | SRR12971560 | - | - | *Marathrum foeniculaceum* (morph) | 11°14'41"N, 73°50'35"W | Caribe/Guachaca |
| AMB 558-2 | SRR12971559 | - | - | *Marathrum foeniculaceum* (morph) | 11°14'41"N, 73°50'35"W | Caribe/Guachaca |
| AMB 558-8 | SRR12971558 | - | - | *Marathrum foeniculaceum* (morph) | 11°14'41"N, 73°50'35"W | Caribe/Guachaca |
| AMB 558-10 | SRR12971557 | - | - | *Marathrum foeniculaceum* (morph) | 11°14'41"N, 73°50'35"W | Caribe/Guachaca |
| AMB 558-11 | SRR12971556 | - | - | *Marathrum foeniculaceum* (morph) | 11°14'41"N, 73°50'35"W | Caribe/Guachaca |
| AMB 562-2 | SRR12971555 | - | - | *Marathrum foeniculaceum* (morph) | 11°13'56"N, 73°42'2"W | Caribe/Don Diego |
| AMB 562-5 | SRR12971554 | - | - | *Marathrum foeniculaceum* (morph) | 11°13'56"N, 73°42'2"W | Caribe/Don Diego |
| AMB 567-3 | SRR12971552 | - | - | *Marathrum foeniculaceum* (morph) | 11°12'2"N, 73°27'42"W | Caribe/Ancho |
| AMB 567-5 | FAILED | - | - | *Marathrum foeniculaceum* (morph) | 11°12'2"N, 73°27'42"W | Caribe/Ancho |
| AMB 567-6 | SRR12971551 | - | - | *Marathrum foeniculaceum* (morph) | 11°12'2"N, 73°27'42"W | Caribe/Ancho |
| AMB 567-7 | SRR12971550 | - | - | *Marathrum foeniculaceum* (morph) | 11°12'2"N, 73°27'42"W | Caribe/Ancho |
| AMB 571-3 | FAILED | - | - | *Marathrum foeniculaceum* (morph) | 11°13'46"N, 73°34'40"W | Caribe/Palomino |
| AMB 571-5 | SRR12971549 | - | - | *Marathrum foeniculaceum* (morph) | 11°13'46"N, 73°34'40"W | Caribe/Palomino |
| AMB 571-6 | SRR12971548 | - | - | *Marathrum foeniculaceum* (morph) | 11°13'46"N, 73°34'40"W | Caribe/Palomino |
| AMB 573-1 | SRR12971547 | - | - | *Marathrum foeniculaceum* (morph) | 11°12'39"N, 73°15'8"W | Caribe/Jerez |
| AMB 573-2 | SRR12971546 | - | - | *Marathrum foeniculaceum* (morph) | 11°12'39"N, 73°15'8"W | Caribe/Jerez |
| AMB 573-3 | SRR12971545 | - | - | *Marathrum foeniculaceum* (morph) | 11°12'39"N, 73°15'8"W | Caribe/Jerez |
| AMB 573-5 | SRR12971544 | - | - | *Marathrum foeniculaceum* (morph) | 11°12'39"N, 73°15'8"W | Caribe/Jerez |
| AMB 573-6 | SRR12971543 | - | - | *Marathrum foeniculaceum* (morph) | 11°12'39"N, 73°15'8"W | Caribe/Jerez |
| AMB 631-2 | SRR12971541 | - | - | *Marathrum foeniculaceum* (morph) | 6°16'11"N, 75°1'54"W | Antioquia/Arenal |
| AMB 631-4 | FAILED | - | - | *Marathrum foeniculaceum* (morph) | 6°16'11"N, 75°1'54"W | Antioquia/Arenal |
| AMB 632 | FAILED | FAILED | - | *Marathrum foeniculaceum* (morph) | 6°16'11"N, 75°1'54"W | Antioquia/Arenal |
| AMB 636 | SRR12971540 | - | - | *Marathrum foeniculaceum* (morph) | 6°16'0"N, 75°1'45"W | Antioquia/Arenal |
| AMB 638-3 | SRR12971539 | - | - | *Marathrum foeniculaceum* (morph) | 6°16'0"N, 75°1'45"W | Antioquia/Arenal |
| AMB 640 | SRR12971538 | - | - | *Marathrum foeniculaceum* (morph) | 6°16'0"N, 75°1'45"W | Antioquia/Arenal |
| AMB 642-3 | FAILED | - | - | *Marathrum foeniculaceum* (morph) | 6°13'57"N, 74°54'58"W | Antioquia/Arenal |
| AMB 645-1 | SRR12971537 | FAILED | - | *Marathrum foeniculaceum* (morph) | 6°11'38"N, 75°0'18"W | Antioquia/Bañadero San Carlos |
| AMB 646-1 | FAILED | FAILED | - | *Marathrum foeniculaceum* (morph) | 6°11'52"N, 75°1'41"W | Antioquia/ |
| AMB 646-4 | SRR12971536 | - | - | *Marathrum foeniculaceum* (morph) | 6°11'52"N, 75°1'41"W | Antioquia/ |
| AMB 646-5 | FAILED | - | - | *Marathrum foeniculaceum* (morph) | 6°11'52"N, 75°1'41"W | Antioquia/ |
| AMB 646-6 | FAILED | - | - | *Marathrum foeniculaceum* (morph) | 6°11'52"N, 75°1'41"W | Antioquia/ |
| AMB 655 | SRR12971535 | - | - | *Marathrum foeniculaceum* (morph) | 5°55'44"N, 75°0'44"W | Antioquia/Samaná Norte |
| AMB 656-1 | SRR12971534 | - | - | *Marathrum foeniculaceum* (morph) | 5°54'56"N, 74°58'45"W | Antioquia/Samaná Norte |
| AMB 658-2 | FAILED | - | - | *Marathrum foeniculaceum* (morph) | 6°0'2"N, 74°56'15"W | Antioquia/Samaná Norte |
| AMB 513 | FAILED | SRR1298324 | - | *Weddellina squamulosa* | 0°51'42.9"N, 71°0'47.5"W | Amazonas/Vaupés |
| AMB 514 | FAILED | - | - | *Weddellina squamulosa* | 0°51'42.9"N, 71°0'47.5"W | Amazonas/Vaupés |
| AMB 515 | FAILED | - | - | *Weddellina squamulosa* | 0°51'42.9"N, 71°0'47.5"W | Amazonas/Vaupés |
| - | - | - | NC_036341 | *Garcinia mangostana* | - | - |
| BNDE | - | - | - | *Hypericum perforatum* | - | - |
| - | - | - | KT337313.1 | *Populus tremula* | - | - |

**Table S2.** Number of loci for which N taxa have data. Ipyrad results for the assembly of all 96 ddRADseq samples and for minimum 4 individuals per loci.

| N | Locus coverage | Sum coverage | N | Locus coverage | Sum coverage | N | Locus coverage | Sum coverage |
| --- | --- | --- | --- | --- | --- | --- | --- | --- |
| 1 | 0 | 0 | 38 | 154 | 168201 | 75 | 0 | 169462 |
| 2 | 0 | 0 | 39 | 136 | 168337 | 76 | 1 | 169463 |
| 3 | 0 | 0 | 40 | 123 | 168460 | 77 | 2 | 169465 |
| 4 | 46519 | 46519 | 41 | 105 | 168565 | 78 | 1 | 169466 |
| 5 | 28200 | 74719 | 42 | 86 | 168651 | 79 | 4 | 169470 |
| 6 | 20962 | 95681 | 43 | 73 | 168724 | 80 | 2 | 169472 |
| 7 | 13485 | 109166 | 44 | 75 | 168799 | 81 | 1 | 169473 |
| 8 | 10892 | 120058 | 45 | 89 | 168888 | 82 | 0 | 169473 |
| 9 | 7606 | 127664 | 46 | 68 | 168956 | 83 | 1 | 169474 |
| 10 | 6563 | 134227 | 47 | 75 | 169031 | 84 | 0 | 169474 |
| 11 | 5050 | 139277 | 48 | 70 | 169101 | 85 | 0 | 169474 |
| 12 | 4442 | 143719 | 49 | 49 | 169150 | 86 | 0 | 169474 |
| 13 | 3471 | 147190 | 50 | 44 | 169194 | 87 | 0 | 169474 |
| 14 | 2973 | 150163 | 51 | 55 | 169249 | 88 | 0 | 169474 |
| 15 | 2310 | 152473 | 52 | 32 | 169281 | 89 | 0 | 169474 |
| 16 | 2112 | 154585 | 53 | 37 | 169318 | 90 | 0 | 169474 |
| 17 | 1721 | 156306 | 54 | 36 | 169354 | 91 | 0 | 169474 |
| 18 | 1510 | 157816 | 55 | 23 | 169377 | 92 | 0 | 169474 |
| 19 | 1210 | 159026 | 56 | 21 | 169398 | 93 | 0 | 169474 |
| 20 | 1131 | 160157 | 57 | 17 | 169415 | 94 | 0 | 169474 |
| 21 | 993 | 161150 | 58 | 16 | 169431 | 95 | 0 | 169474 |
| 22 | 868 | 162018 | 59 | 6 | 169437 | 96 | 0 | 169474 |
| 23 | 762 | 162780 | 60 | 8 | 169445 |  |  |  |
| 24 | 663 | 163443 | 61 | 2 | 169447 |  |  |  |
| 25 | 563 | 164006 | 62 | 1 | 169448 |  |  |  |
| 26 | 555 | 164561 | 63 | 1 | 169449 |  |  |  |
| 27 | 454 | 165015 | 64 | 1 | 169450 |  |  |  |
| 28 | 499 | 165514 | 65 | 2 | 169452 |  |  |  |
| 29 | 391 | 165905 | 66 | 4 | 169456 |  |  |  |
| 30 | 375 | 166280 | 67 | 0 | 169456 |  |  |  |
| 31 | 349 | 166629 | 68 | 0 | 169456 |  |  |  |
| 32 | 314 | 166943 | 69 | 1 | 169457 |  |  |  |
| 33 | 254 | 167197 | 70 | 0 | 169457 |  |  |  |
| 34 | 245 | 167442 | 71 | 1 | 169458 |  |  |  |
| 35 | 202 | 167644 | 72 | 1 | 169459 |  |  |  |
| 36 | 211 | 167855 | 73 | 3 | 169462 |  |  |  |
| 37 | 192 | 168047 | 74 | 0 | 169462 |  |  |  |

**Table S3.**
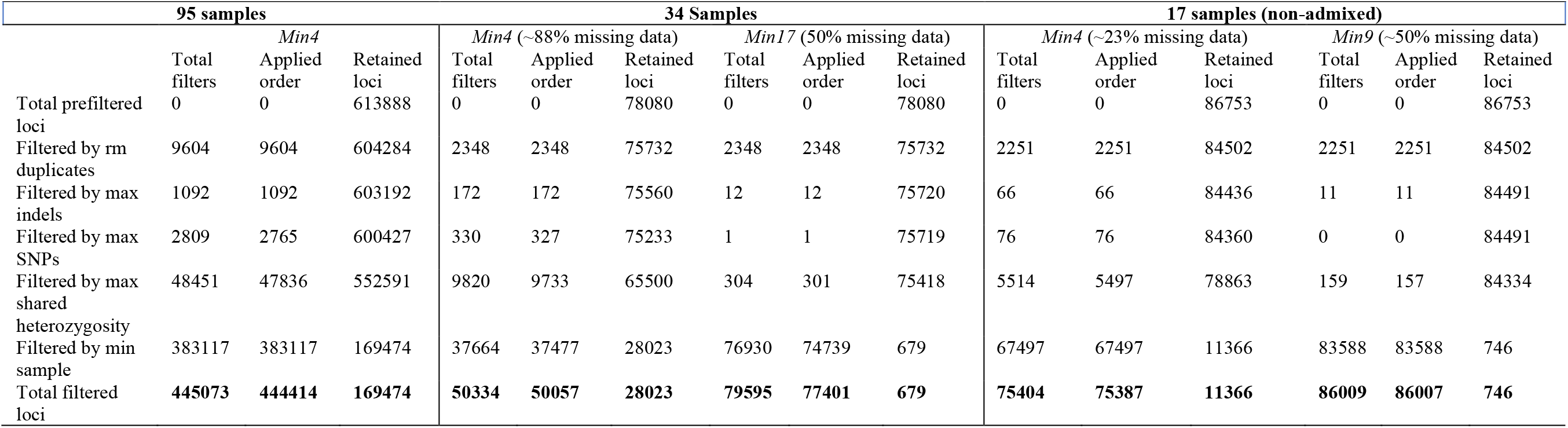
Ipyrad assembly statistics for our total 95, subset of 34 admixed and non-admixed, and 17 non-admixed ddRADseq samples. The latter two were also included in target enrichment. Statistics are shown for loci shared across minimum 4 and 17 of the samples in our subset of 34 samples, and minimum 4 and 9 samples in our subset of 17 non-admixed individuals.

**Table S4.** Target enrichment recovery efficiency. The table includes the number of sequenced reads and of exons recovered out of the total 536 targeted. Target length (TL). The table is organized in descending order from the sample with most loci recovered at 75% of the TL. *Marathrum* samples with relatively short exons were excluded from downstream analyses but the outgroup was included. The first sample listed is the one used for selection of sequences to be targeted with the pipeline Sondovac. Admixed individuals with the phenotype of *M. foeniculaceum* are indicates with quotation marks.

| Sample | # Reads | # Exons recovered |  |  | Loci recovered |
| --- | --- | --- | --- | --- | --- |
|  |  | >25% TL | >50% TL | >75% TL |  |
| *M. utile* AMB497 | 18152507 | 535 | 535 | 534 | 208 |
| *M. utile* AMB574-1 | 6015552 | 529 | 527 | 527 | 208 |
| *M. utile* AMB503 | 20726495 | 526 | 526 | 526 | 208 |
| *M. foeniculaceum* AMB643-1 | 4444374 | 526 | 526 | 526 | 208 |
| *M. foeniculaceum* AMB571-1 | 6308716 | 527 | 526 | 526 | 208 |
| *M. utile* AMB659 | 6645871 | 526 | 526 | 526 | 208 |
| *M. utile* AMB641-1 | 6598011 | 526 | 525 | 524 | 208 |
| *M. utile* AMB653 | 11275084 | 523 | 522 | 522 | 208 |
| *M. utile* AMB512 | 4921124 | 525 | 523 | 522 | 208 |
| *M. utile* AMB652 | 10640199 | 523 | 522 | 521 | 208 |
| "*M. foeniculaceum*" AMB494 | 8038995 | 529 | 527 | 521 | 203 |
| "*M. foeniculaceum*" AMB499 | 9799316 | 531 | 529 | 521 | 208 |
| *M. foeniculaceum* AMB567-4 | 845956 | 524 | 522 | 520 | 208 |
| *M. utile* AMB639-1 | 4951653 | 521 | 521 | 519 | 208 |
| *M. utile* AMB507 | 843464 | 519 | 519 | 519 | 208 |
| *M. utile* AMB577-3 | 9935840 | 520 | 519 | 519 | 208 |
| *M. foeniculaceum* AMB635 | 4098778 | 522 | 520 | 518 | 208 |
| "*M. foeniculaceum*" AMB562-1 | 871580 | 522 | 521 | 518 | 202 |
| *M. utile* AMB577-2 | 15127071 | 519 | 518 | 517 | 208 |
| "*M. foeniculaceum*" AMB571-4 | 6618910 | 530 | 527 | 517 | 203 |
| "*M. foeniculaceum*" AMB502 | 11814928 | 526 | 524 | 516 | 203 |
| "*M. foeniculaceum*" AMB561-2 | 2350936 | 528 | 526 | 516 | 203 |
| *M. utile* AMB575-1 | 1971791 | 517 | 517 | 516 | 208 |
| "*M. foeniculaceum*" AMB654 | 10076885 | 526 | 524 | 515 | 203 |
| *M. utile* AMB578 | 9519370 | 516 | 516 | 515 | 208 |
| *M. utile* AMB566-4 | 1589623 | 518 | 517 | 515 | 208 |
| "*M. foeniculaceum*" AMB558-5 | 3386292 | 528 | 523 | 513 | 203 |
| "*M. foeniculaceum*" AMB658-1 | 4293245 | 521 | 517 | 513 | 203 |
| "*M. foeniculaceum*" AMB573-4 | 8038288 | 525 | 524 | 512 | 203 |
| "*M. foeniculaceum*" AMB638-4 | 1878381 | 521 | 518 | 512 | 203 |
| "*M. foeniculaceum*" AMB558-4 | 2851635 | 525 | 522 | 512 | 203 |
| "*M. foeniculaceum*" AMB561-3 | 591426 | 514 | 513 | 510 | 203 |
| "*M. foeniculaceum*" AMB657 | 2993130 | 520 | 516 | 510 | 203 |
| "*M. foeniculaceum*" AMB506 | 11255969 | 518 | 513 | 507 | 203 |
| "*M. foeniculaceum*" VLondono | 7047222 | 515 | 512 | 506 | 208 |
| "*M. foeniculaceum*" AMB555-9 | 507428 | 506 | 506 | 503 | 203 |
| "*M. foeniculaceum*" AMB551-3 | 10403228 | 521 | 514 | 500 | 208 |
| *M. foeniculaceum* AMB571-2 | 1658509 | 517 | 506 | 479 | 208 |
| "*M. foeniculaceum*" AMB568 | 576529 | 498 | 492 | 478 | 202 |
| *M. utile* AMB563-1 | 497235 | 489 | 484 | 470 | 206 |
| "*M. foeniculaceum*" AMB556-2 | 447726 | 445 | 431 | 405 | 201 |
| ***M. foeniculaceum* AMB646-1** | 528950 | 481 | 439 | 341 | -- |
| ***M. utile* AMB642-1** | 269227 | 450 | 387 | 263 | -- |
| *M. foeniculaceum* AMB632 | 125422 | 305 | 204 | 119 | -- |
| *M. foeniculaceum* AMB645 | 364733 | 90 | 60 | 47 | -- |
| *W. squamulosa* AMB513 | 893791 | 318 | 285 | 232 | 203 |

**Figure S1.**
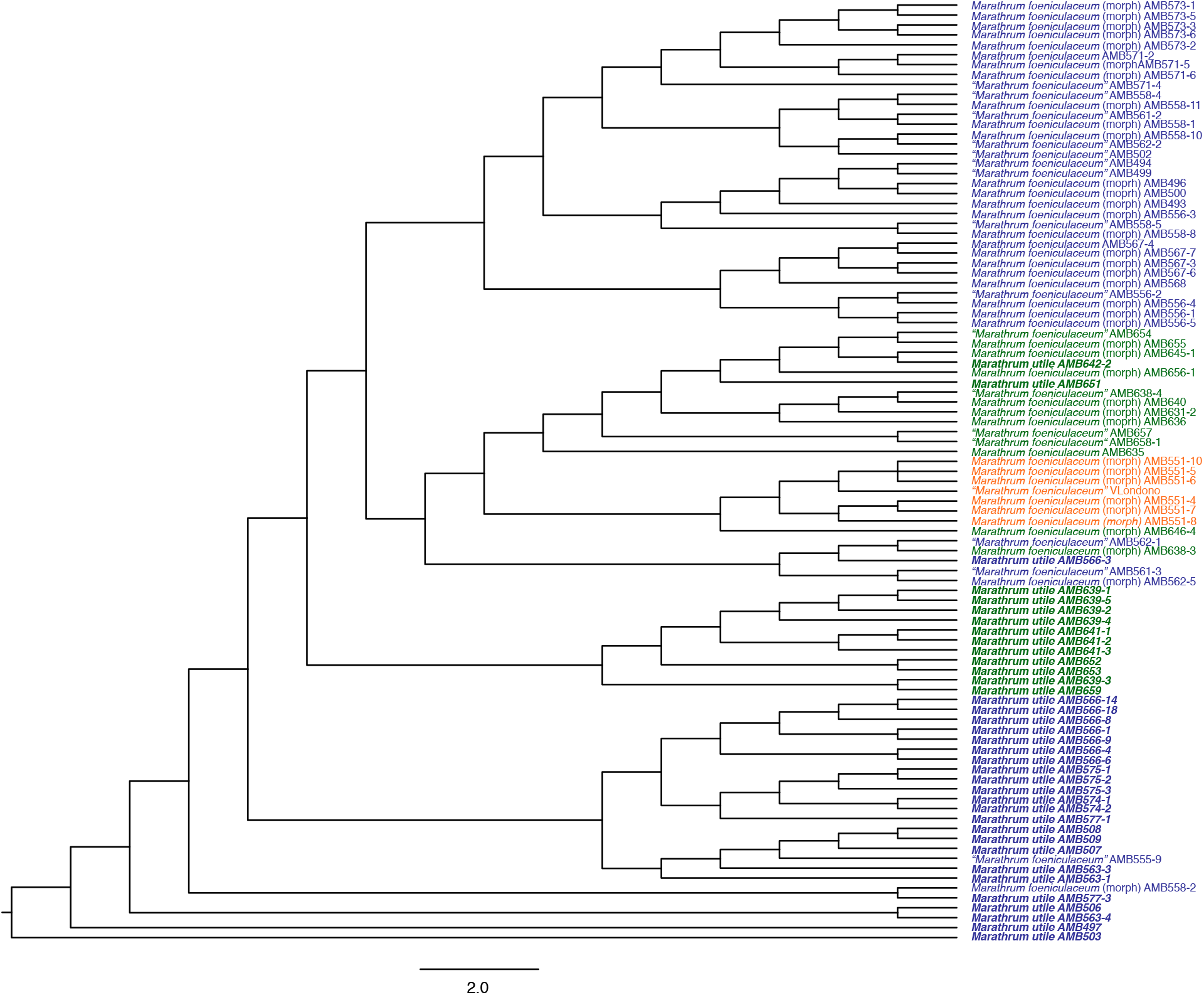
Species tree inferred with SVDQuartets for 95 ddRADseq samples successfully sequenced in this study. Individuals of *M. foeniculaceum* and *M. utile* (bold) from Antioquia (green), Boyacá (orange) and Caribe (blue) are shown. Tree was inferred as unrooted, but rotting is presented to be consistent with posterior analyses that were rooted with the outgroup *W. squamulosa*. Admixed individuals as inferred with ADMIXTURE are indicated with quotation marks. Individuals for which no information on genetic constitution is available but have the phenotype of *M. foeniculaceum* are indicated with “(morph)”.

**Figure S2.**
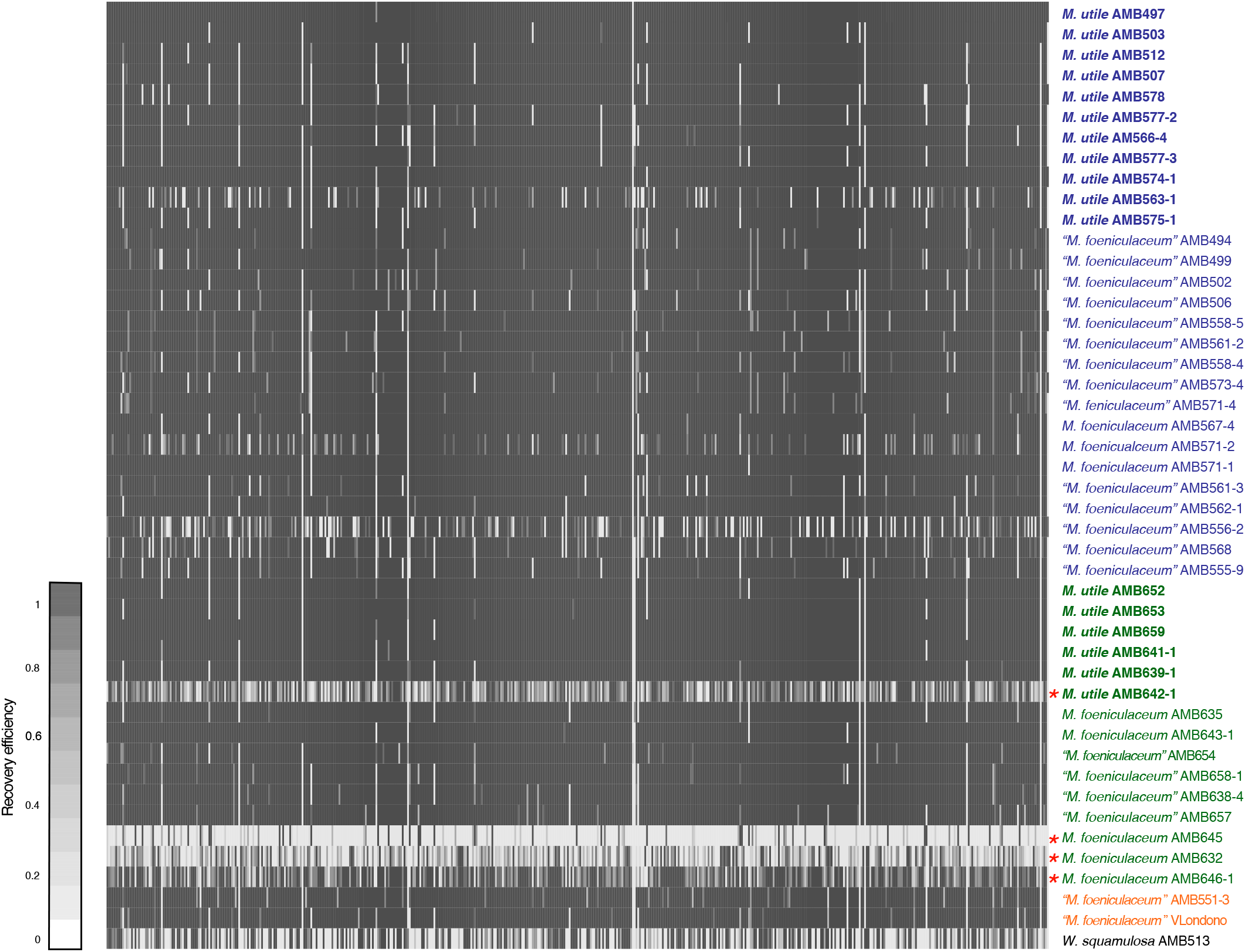
Target enrichment loci recovery efficiency for 42 samples (rows) across loci (columns). Recovery efficiency is measured as the length of the sequence recovered as a percentage of the length of the targeted gene. Darker gray: The full length of the targeted sequence was recovered. White: no sequence was recovered. (*) Individuals that failed sequencing or recovered short exons of low quality. Individuals of *M. foeniculaceum* and *M. utile* (bold) from Antioquia (green), Boyacá (orange) and Caribe (blue) are shown. Admixed individuals as inferred with ADMIXTURE are indicated with quotation marks.

**Figure S3.**
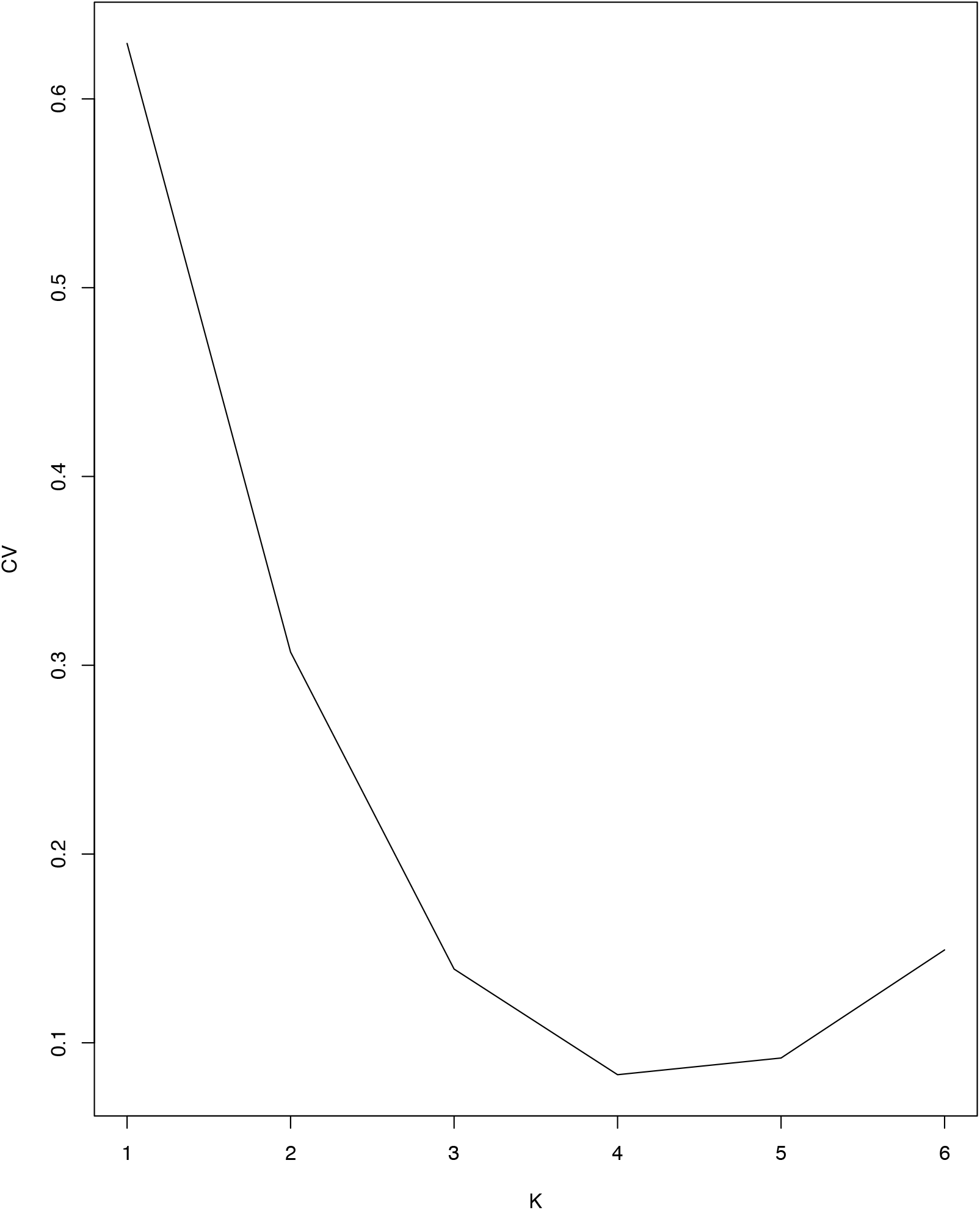
Cross-validation error and number of clusters (K) resulted for population genetics analysis with ADMIXTURE using target enrichment data.

**Figure S4.**
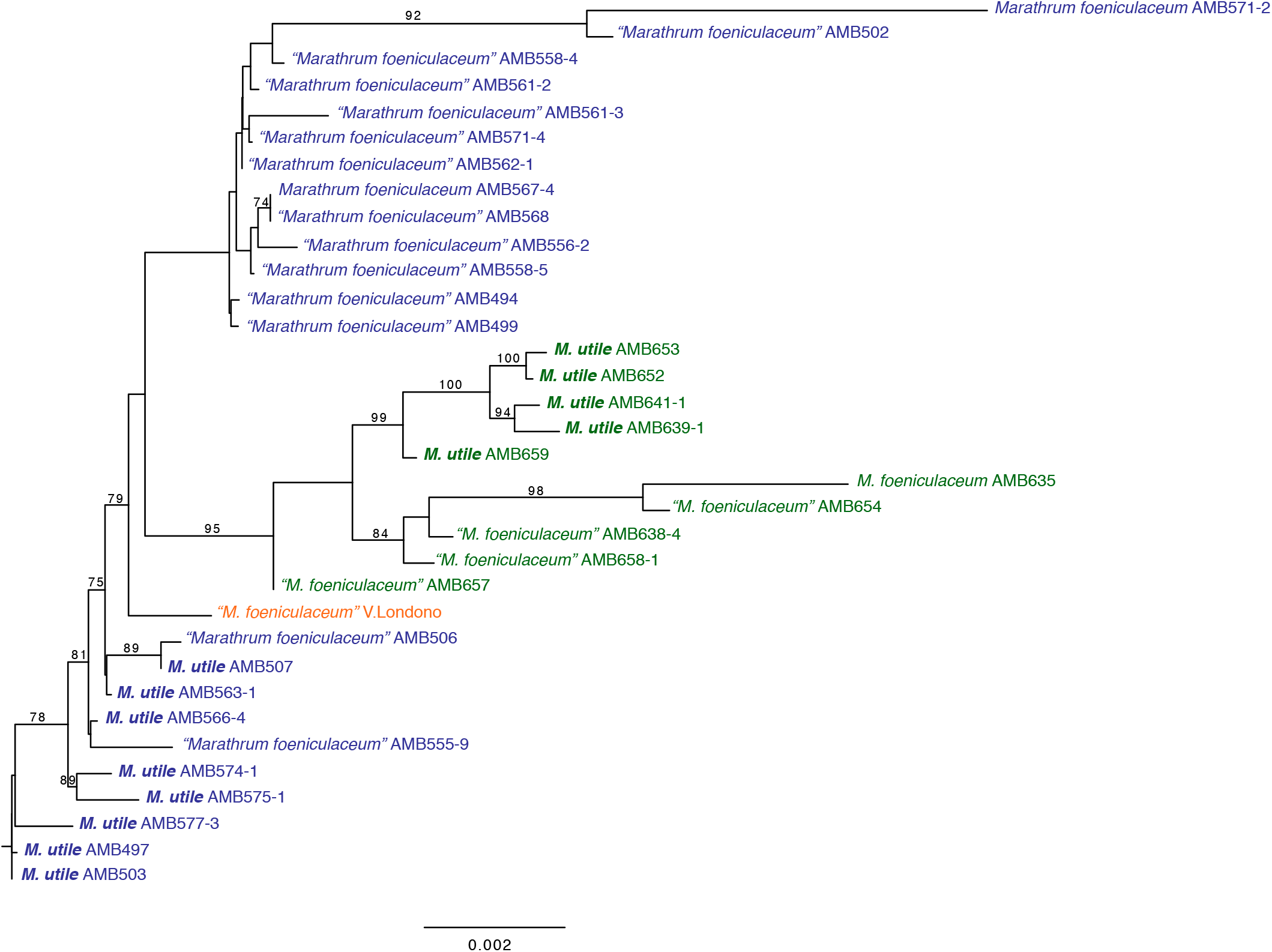
Maximum likelihood tree inferred from our ddRADseq dataset including only individuals that are also in the target enrichment dataset. Individuals of *M. foeniculaceum* and *M. utile* (bold) from Antioquia (green), Boyacá (orange) and Caribe (blue) are shown. Admixed individuals as inferred with ADMIXTURE are indicated with quotation marks.

**Figure S5.**
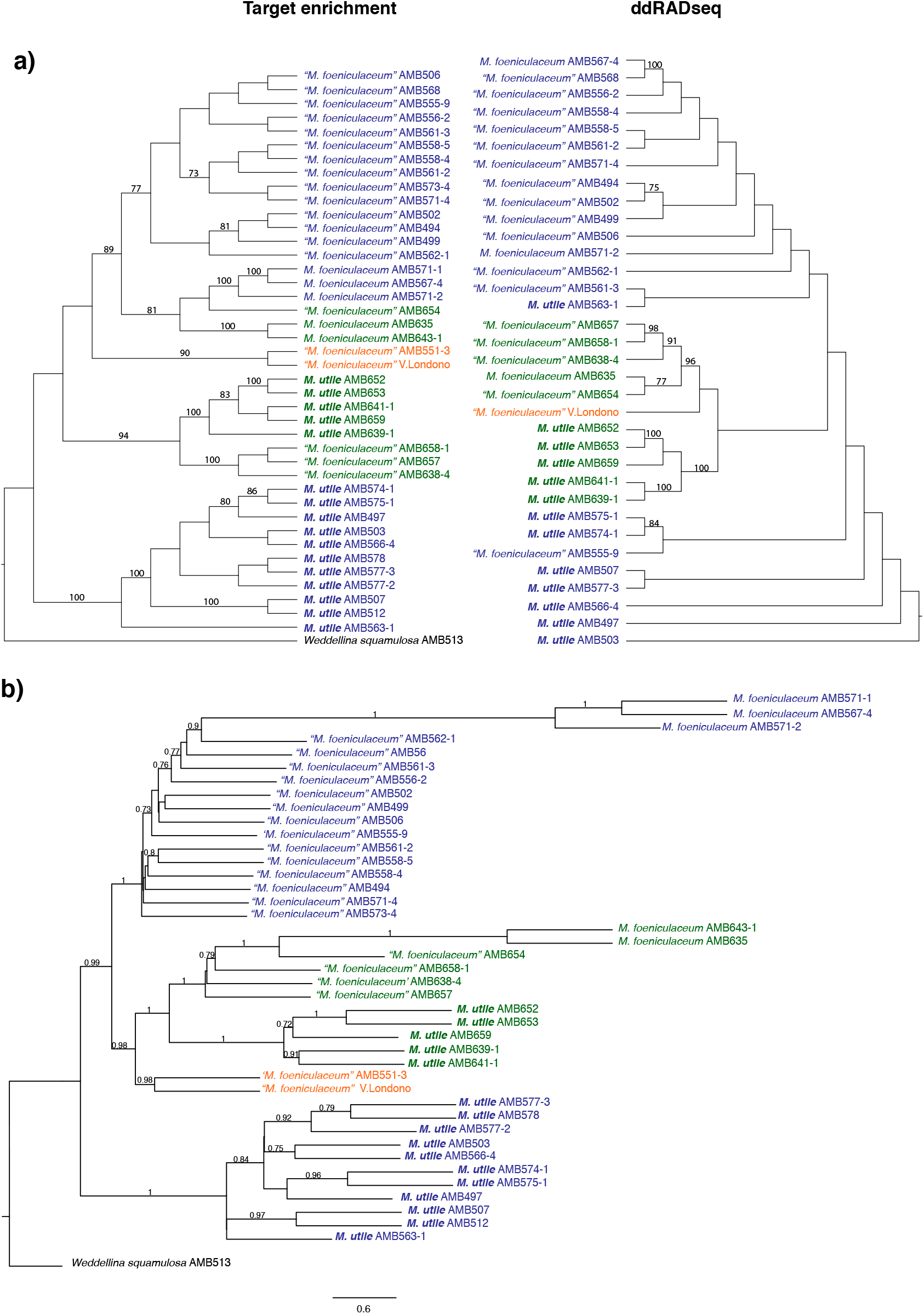
Species trees inferred from our target enrichment and ddRADseq data. a) SVDQuartets tree with quartet support values >70 above branches and b) ASTRAL III tree with local posterior probabilities > 0.7 shown above branches. Individuals of *M. foeniculaceum* and *M. utile* (bold) from Antioquia (green), Boyacá (orange) and Caribe (blue) are shown. Admixed individuals as inferred with ADMIXTURE are indicated with quotation marks.

**Figure S6.**
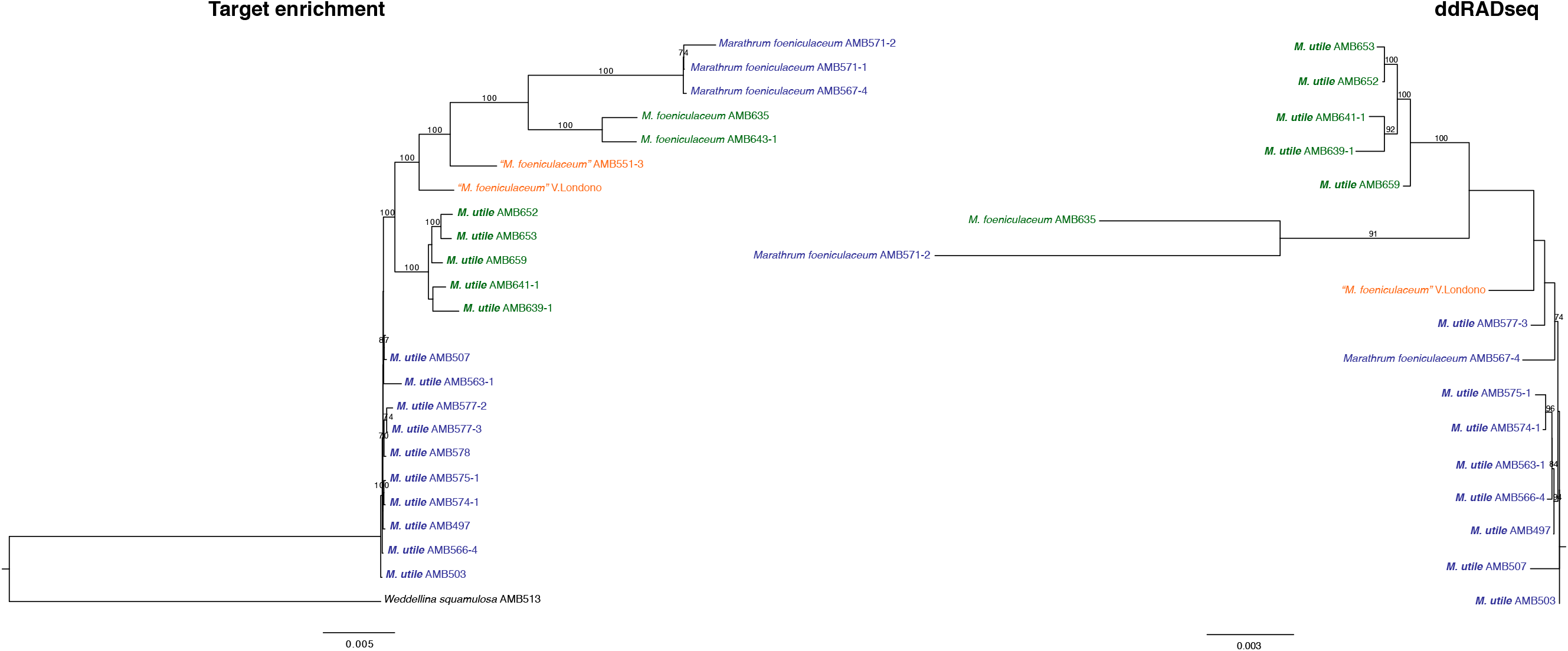
Maximum likelihood trees inferred from our target enrichment and ddRADseq data excluding admixed individuals from Antioquia and Caribe. Bootstrap support values >70 are show above branches. Individuals of *M. foeniculaceum* and *M. utile* (bold) from Antioquia (green), Boyacá (orange) and Caribe (blue) are shown. Admixed individuals from Boyacá are indicated with quotation marks.

**Figure S7.**
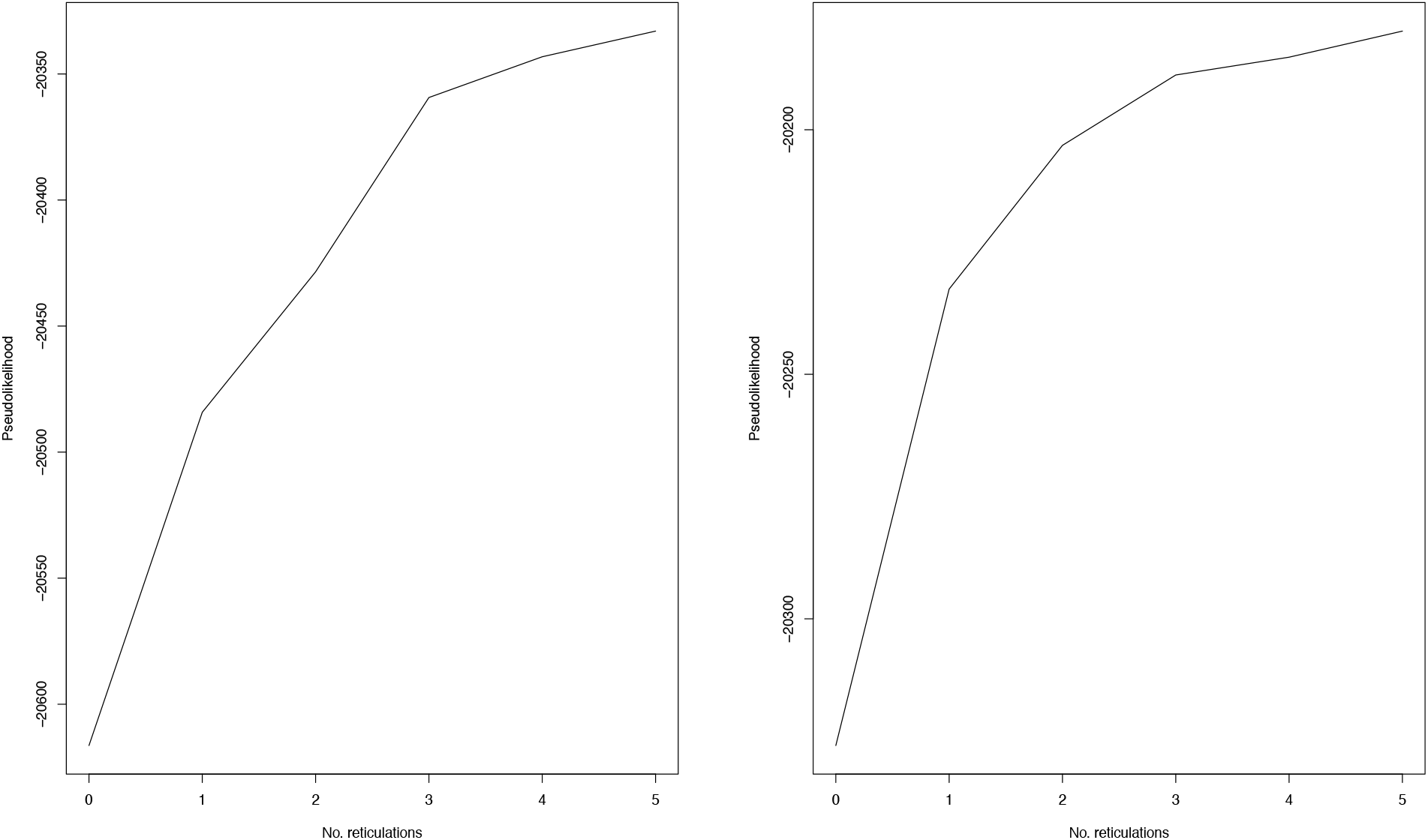
PhyloNet results allowing 0–5 reticulations and including admixed individuals. After 5 reticulation events, the change in pseudolikelihood score has not plateaued. Each graph represents a separate dataset with 2 semi-randomly chosen individuals per population (Antioquia, Boyacá and Caribe).

